# Disinhibition of Plant Defense Responses as an Innovative Biocontrol Strategy in Grapevine

**DOI:** 10.64898/2026.11.19.767271

**Authors:** Chloé Ilbert, Jérémy Villette, Sébastien Aimé, Manon Chargy, Marie-Cécile Dufour, Caroline Bonneau, Cécile Blanchard, Karine Palavioux, Hoai-Nam Truong-Cellier, Nathalie Harzic, Marielle Adrian, Stéphane Bourque

## Abstract

Grapevine (*Vitis vinifera*) is susceptible to numerous fungal and oomycete pathogens. Plant defense inducers (PDIs) offer the potential to reduce the use of chemical fungicides in viticulture, but their efficacies under vineyard conditions remain variable. Here, we investigated whether alleviating the repression of immune responses could enhance PDI-induced protection against grapevine diseases. We first characterized two grapevine HD2-type histone deacetylases, VvHD2a and VvHD2b, as negative regulators of basal immunity. Their constitutive expression in *Arabidopsis thaliana* attenuated the flg22-induced oxidative burst and defense-related gene expression, while increasing susceptibility to *Botrytis cinerea*. The histone deacetylase inhibitor, sodium butyrate (NaBut), blocked this enhanced susceptibility. In grapevine, NaBut alone protected against *B. cinerea* but did not against *Plasmopara viticola*. However, combining NaBut with low doses of the PDI Belvine^®^ conferred protection against *P. viticola* comparable to that achieved with the full recommended PDI dose, with a synergistic interaction between both treatments. Similarly, NaBut enhanced Belvine^®^-induced signaling events, such as MAPK phosphorylation and transcriptomic analysis revealed amplified expression of genes involved in multiple defense-related processes. Finally, field trials conducted in two Bourgogne vineyards showed that adding NaBut to a reduced-copper program incorporating Belvine^®^ improved downy mildew control on leaves and bunches, with Bayesian probabilities of improved protection ranging from 91% to 95%. Disease severity was comparable to that achieved with the full copper dosage. These findings highlight the potential of combining defense induction by a reduced dosage of PDI and disinhibition to improve PDI-induced disease control while reducing copper inputs in viticulture.

**Highlights:**

- Sodium butyrate-mediated disinhibition of grapevine defenses enhances Belvine^®^-induced protection against *Plasmopara viticola*.
- Biochemical and transcriptomic analyses highlight how sodium butyrate modulates Belvine^®^-induced defense responses.
- Field trials validate the protective effect of the disinhibition strategy against *P. viticola* on grape leaves and clusters.
- Combining defense induction with sodium butyrate-mediated disinhibition represents a promising strategy to strengthen high level in grapevine protection while reducing fungicides.

## 1. Introduction

Faced with the effects of climate change and growing environmental and societal expectations, viticulture must adapt its practices and move toward more sustainable production strategies (Bisson et al., 2002). However, this remains a true challenge because grapevine (*Vitis vinifera*) is highly susceptible to a wide range of pests and pathogens, including insects, viruses, oomycetes such as *Plasmopara viticola* (responsible for downy mildew), and fungus like *Erysiphe necator* (powdery mildew) or *Botrytis cinerea* (grey mold). Among the different diseases of grapevine, downy (DM) and powdery (PM) mildews are the most damaging in vineyards causing substantial yield losses and affecting berry and wine quality. Disease control still relies largely on repeated chemical treatments, which raise growing concerns for the environment and human health (De Bernardi et al., 2022; Komárek et al., 2010; Pathak et al., 2022). These issues have encouraged the development of alternative protection strategies. These include approaches based on natural mechanisms, such as the use of antagonistic microorganisms, natural extracts, or plant defense inducers (PDIs). While the two first strategies act directly on pathogens, PDIs stand out for their ability to stimulate the plant’s immune system and enhance its resistance to pathogens (Adrian et al., 2017, 2024; Walters et al., 2013). Despite recent advances, the effectiveness of PDIs in vineyards often remains variable and frequently limited, limiting their large-scale adoption (Brulé et al., 2024; Dagostin et al., 2011; Fedele et al., 2020). For instance, already existing homologated biocontrol products in Europe such as COS-OGA-based products (containing chitooligosaccharides and oligogalacturonides) or yeast extracts provide good levels of protection against oomycetes and fungal pathogens under greenhouse conditions (Goncharuk et al., 2022). However, when applied in vineyards, their efficacy can still be variable (Dagostin et al., 2011; Fedele et al., 2020; Brulé et al., 2024). These PDIs are mainly composed of molecular motifs that can be recognized by plant cell-surface receptors. These motifs, called MAMPs (Microbe-Associated Molecular Patterns) or DAMPs (Damage-Associated Molecular Patterns), are perceived by pattern-recognition receptors (PRRs) located at the plasma membrane (Ngou et al., 2022). Their recognition activates a first layer of immunity, known as PTI (Pattern-Triggered Immunity), which initiates a complex signalling cascade response. However, evolution selected pathogens that succeeded in circumventing this first immune barrier by secreting virulence molecules, known as effectors, which are able to neutralize basal host defences. Then, plants that had developed a second level of immunity, called ETI (Effector-Triggered Immunity) were selected, based on the direct or indirect recognition of these effectors by intracellular NB-LRR (Nucleotide-Binding Leucine-Rich Repeat) proteins. ETI triggers a hypersensitive response (HR; Jones and Dangl, 2006). In plants, both PTI and ETI activate interconnected signalling networks that lead to a cascade of signalling events such as the production of Reactive Oxygen Species (ROS), the activation of Mitogen-Activated Protein Kinases (MAPKs) and subsequently the activation of defence gene expression followed by the establishment of effective defence responses (Meng and Zhang, 2013; Ngou et al., 2021; Yuan et al., 2021). These immune responses are finely regulated by phytohormonal pathways, notably involving salicylic acid (SA), jasmonic acid (JA), and ethylene (ET), which coordinate the amplitude and specificity of downstream defence mechanisms (Huang et al., 2026; Kelloniemi et al., 2015; Tsuda and Katagiri, 2010). A hallmark of grapevine immunity is the production of stilbene phytoalexins, particularly resveratrol and its oligomeric derivatives such as viniferins, which accumulate upon pathogen challenge (Jeandet et al., 2002; Chong et al., 2009). These compounds represent a major component of the inducible chemical defence arsenal in *V. vinifera*, contributing to the restriction of pathogen spread and the reinforcement of host resistance.

In the context of plant immunity, epigenetic regulation has been shown to modulate defence responses to a wide range of PDIs and pathogens (Hannan Parker et al., 2022). Among the different types of epigenetic modifications, histone acetylation plays a central role in the regulation of these responses (Ramirez-Prado et al., 2018). Histone acetylation dynamics are primarily controlled by the antagonistic activities of Histone Acetyltranferases (HAT) and Histone Deacetylases (HDACs), which regulate the acetylation levels of specific lysine residues on both histone and non-histone proteins (Villette et al., 2025). HDACs are key components of plant immunity, with several members reported to act as either positive or negative regulators of resistance to pathogens (An et al., 2022; Bourque et al., 2011; Choi et al., 2012; Ding et al., 2012; Kim et al., 2008; Wang et al., 2017; Zhou et al., 2005). Notably, plants possess a specific HDAC subfamily, the HD2 proteins (type-2 HDACs), which form a distinct class separated from the other two HDAC subfamilies conserved across eukaryotes (Bourque et al., 2016; Grandperret et al., 2014). Functional studies have shown that HD2 proteins in tobacco and rice act as negative regulators of plant immunity (Bourque et al., 2011; Ding et al., 2012). For instance, in rice, HDT701 has been shown to repress defence activation and reduce resistance to pathogens (Ding et al., 2012).

To alleviate this repression of plant immunity by HDACs, approaches involving the use of HDAC inhibitors have been developed. Indeed, hundreds of HDAC inhibitors have been identified, primarily through cancer research, including sodium butyrate (NaBut; Anh et al., 2021; Davie, 2003). NaBut is a short-chain fatty acid naturally present in various biological environments and widely used in industrial applications such as food production, animal feed, pharmaceutics and cosmetics (Coppola et al., 2022; Gerunova et al., 2024). In several model systems, particularly in animals and microorganisms, NaBut and its derivatives have been shown to inhibit HDAC activity, leading to increased histone acetylation and the induction of genes involved in cell protection and immune responses (Steliou et al., 2012; Xiong et al., 2016). In plants, recent studies have demonstrated the effects of NaBut in rice, where its application increases histone acetylation levels, including H3K9 and H4K5 acetylation, consistent with the inhibition of HDAC activity. NaBut treatment also induces defense-related genes and enhances resistance to rice blast disease (Xu et al., 2022). This raises the question of whether the effect of NaBut on plant immunity may involve the inhibition of HD2 proteins. Interestingly, silencing of HD2 genes in rice and tobacco has also been associated with enhanced defense responses and resistance (Bourque et al., 2011; Ding et al., 2012). Altogether, these findings suggest that NaBut can relieve HDAC-mediated transcriptional repression and raise the possibility that HD2 proteins may contribute to this effect, thereby creating a chromatin state that may facilitate the activation of defense-related genes upon subsequent stimulation.

In the present study, we investigated the role of grapevine HD2 proteins in plant immunity using constitutive expression in the model plant *Arabidopsis thaliana*. Our results show that grapevine HD2s act as negative regulators of immunity during fungal infection, leading us to hypothesize that their inhibition by NaBut could relieve transcriptional constraints on grapevine defense responses. We therefore used various approaches to investigate whether NaBut could exert a protective effect against pathogen infection on its own. We then explored whether NaBut-mediated disinhibition could further enhance grapevine responsiveness to a PDI. Overall, our results support a role for NaBut in relieving HDAC mediated defense repression in grapevine and thereby enhancing the PDI effectiveness against DM, both under controlled and field conditions.

## 2. Materials and methods

### 2.1. Plant materials and growth condition

#### 2.1.1. Arabidopsis thaliana

*A. thaliana* Columbia ecotype (Col-0-WT) and transgenic lines were grown at 20/18°C under a 10 h light/14 h dark photoperiod. Transgenic plants expressing either VvHD2a (Vitvi06g01399) or VvHD2b (Vitvi08g01518) were generated using the floral-dip method (Clough and Bent, 1998). The coding sequences of VvHD2a and VvHD2b, amplified from *V. vinifera* cv. Chardonnay leaf cDNA, were cloned into the pFAST_R02 vector for constitutive expression and pFAST_R05 for constitutive expression with a Green Fluorescent Protein (GFP) tagged in N-terminal of each VvHD2. For each construct, two independent homozygous lines were established.

#### 2.1.2. Vitis vinifera

Herbaceous cuttings of *V. vinifera* cv. Marselan were planted individually in 8×8×8 cm pots containing a blond peat-perlite mixture (4:1, v/v). Plants were cultivated in a glasshouse at 24°C during the day and 18°C at night, with a 16-hour photoperiod and relative humidity maintained at 70 ± 10%. Irrigation was provided through a drip system using a balanced NPK fertilizer solution (10-10-10; Plantin, France). Experiments were conducted seven weeks after rooting, when plants had developed six to eight fully expanded leaves.

*In vitro* plantlets of *V. vinifera* cv. Marselan were cultivated in glass tubes (2 cm diameter, 15 cm height) containing 15 mL of culture medium (Table S1) solidified with 7.5 g.L⁻¹ agar (Duchefa, P1003), and subcultured every two months. Cultures were maintained in a growth chamber at 24 ± 2°C under cool-white fluorescent lighting (3350 L) with a 16 h light / 8 h dark photoperiod.

### 2.2. Treatments with NaBut and PDIs

For the induction of plant defense responses in *A. thaliana*, the flg22 peptide from *Xanthomonas campestris* pv. *campestris* strain 305 (QRLSSGLRINSAKDDAAGLAIS) was used as a PDI at a final concentration of 1 µM in ultrapure water. For RT-qPCR experiments, leaves from four-week-old *A. thaliana* plants were pre-infiltrated with ultrapure water and placed in six-well plates containing ultrapure water for 4 h. The water was then replaced with either ultrapure water (control) or flg22 solution, and leaves were incubated for 1 or 3 h before being collected and immediately frozen in liquid nitrogen. For ROS assays, two leaf discs from four-week-old *A. thaliana* plants were collected from two independent plants and placed in separated wells, with each representing one replicate. NaBut treatments were performed using NaBut with purity ≥ 99% (Sigma-Aldrich, ref. 303410), freshly prepared in ultrapure water to a final concentration of 1 mM. Four-week-old *A. thaliana* plants were treated by foliar spraying 24 h prior to *B. cinerea* or *Pseudomonas syringae pv. tomato DC3000* infection.

For grapevine assays, a formulated commercial PDI produced and supplied by Cérience (Belvine^®^, France), was used at a reduced dose compared with its recommended field application rate (3.25 g.L⁻¹), corresponding to 0.75 g.L⁻¹ in all experiments, alone or in combination with NaBut (1 mM). For combined treatments, plants were first treated with NaBut, followed by the application of Belvine^®^ 3 h later. This interval was chosen to allow the first treatment to dry on the leaf surface before applying the second treatment. All solutions were prepared in sterile ultrapure water and applied immediately after preparation. Tests were performed on treated grapevine plants to evaluate the protection induced by NaBut and Belvine^®^ against *P. viticola* and *B. cinerea.* Six plants per treatment and condition were used. On day 0, the first fully expanded leaves (ranks 1 and 2 from the apex) were sprayed on both sides with the treatment solutions, ensuring a fine and homogeneous mist without runoff. Plants were left to dry for at least 3 h and then grown in the glasshouse under irrigation. Pathogen inoculation was performed 48 h later.

### 2.3. Pathogen infection assays

#### 2.3.1. For Arabidopsis thaliana

The *B. cinerea* inoculum was prepared by culturing the BMM strain (Zimmerli et al., 2001) on a solid V8 medium (50% V8 tomato juice in ultrapure water – v/v – supplemented with 36 mM KH_2_PO_4_ and 3% agar, adjusted to pH = 6) in the dark to promote sporulation. Conidia were then harvested, concentrated in water, and diluted to a final concentration of 5 × 10⁴ conidia.mL^-1^ using 0.6% potato dextrose broth (PDB) as the diluent. Four-week-old *A. thaliana* leaves were inoculated with 6 µL droplets of the conidial suspension, corresponding to approximately 300 conidia per leaf. Lesion diameters were measured three days post-inoculation to assess necrosis. Based on the protocol described in Nguyen et al. (2023), *P. syringae* pv. *tomato* DC3000 was cultured at 28°C. A bacterial glycerol stock was streaked onto King’s B agar plates containing 20 g.L⁻¹ peptone special (Merck Millipore, Germany), 1% (v/v) glycerol, 11 mM KH_2_PO_4_, 6 mM MgSO_4_-7H_2_O, and 15 g.L⁻¹ agar, adjusted to pH 7.2, and supplemented with 50 µg.mL⁻¹ rifampicin. After two days of incubation, a single colony was picked and grown overnight in King’s B liquid medium with constant shaking. Fifty microliters of this overnight culture were then inoculated into 50 mL of King’s B liquid medium and incubated overnight with continuous shaking to reach an optical density of 0.2 at 600 nm (OD₆₀₀). Cells were harvested by centrifugation (3000 × g, 15 min) and resuspended in 10 mM MgCl_2_. Serial dilutions in 10 mM MgCl_2_ were performed to reach 10⁶ CFU.mL⁻¹.

For each Arabidopsis genotype, three leaves from three 4-week-old plants were infiltrated with the bacterial suspension using a 1 mL needleless syringe. Plants were maintained in a tray covered with a transparent lid to create a high-humidity environment. After 24 h, the lid was removed. Forty-eight hours post-infiltration, 7 mm diameter leaf discs were collected from each infiltrated leaf for each genotype, ground with a mortar and pestle, and resuspended in 10 mL of 10 mM MgCl_2_ to extract the bacteria. Finally, 150 µL of tenfold serial dilutions (10⁻² to 10⁻⁵) were plated onto King’s B agar plates and incubated at 28°C. Bacterial colonies were counted after two days.

#### 2.3.2. For Vitis vinifera

The DM pathogen *P. viticola* (originally collected in a Burgundy vineyard, SRPV Beaune, France) was routinely maintained on *V. vinifera* cv. Marselan plants grown under the glasshouse conditions described above. For inoculation, sporangia were collected in distilled water from a freshly sporulating leaf. The suspension was adjusted to 1 × 10⁴ sporangia.mL⁻¹ and sprayed onto the abaxial leaf surface to produce a thin, homogeneous film of droplets. After inoculation, plants were kept in darkness at 100% relative humidity for at least 6 h to promote zoospore release and infection, then grown in greenhouses. Five days later, disks (1.2 cm diameter) were excised from leaves, placed on moistened Whatman paper, and incubated at 100% humidity to induce sporulation under a 10 h/14 h light/dark cycle at 20/18°C. Infection intensity was assessed at 6, 7 and 8 days post-inoculation (dpi) by imaging disks and quantifying the sporulating area using Visilog 6.9 image analysis software (Kim Khiook et al., 2013).

For *B. cinerea* infection assays, 40 leaf disks (2 cm diameter) from two leaves (ranks 1 and 2) of four plants were collected, placed on moistened Whatman paper, and inoculated with a 20 µL droplet of the conidial suspension at a concentration of 5 × 10⁴ conidia mL⁻¹ in 0.6% (w/v) of PDB. Disks were incubated at 100% humidity under a 14 h/10 h light/dark cycle at 20/18°C. Lesion development was monitored 3 dpi by photographing disks and measuring lesion diameter using the ImageJ software.

### 2.4. Evaluation of the direct effect

The direct effect of treatments on *P. viticola* was evaluated through zoospore mobility assay. Sporangia were collected from freshly sporulating grapevine leaves and suspended in sterile water to obtain a final concentration of 1 × 10⁵ sporangia.mL⁻¹. Zoospore release was first allowed to occur in water for 1-2 h. The following treatments were tested: sterile water (control), NaBut at four concentrations (0.1, 1, 5 and 10 mM), Belvine^®^ at two concentrations (0.75 and 3.25 g·L⁻¹), and the combination of NaBut (1 mM) and Belvine^®^ (0.75 g·L⁻¹). All concentrations refer to final concentrations in the reaction mixture. For each treatment, 200 µL were then added to 1.8 mL of the zoospore suspension. Two minutes after treatment application, a 20 µL aliquot was collected and loaded into a Malassez counting chamber for mobility assessment. Videos (1 min in duration) were recorded at low magnification using a microscope-integrated camera to capture several grid squares simultaneously (BRESSER researcher trino 40-1000x microscope, 14334). For each treatment, zoospore mobility was quantified from three independent videos, and in each video the number of zoospores entering three distinct grid squares per minute was recorded (Video S1-S8). Mobility was also re-evaluated at later time points (up to 1 h) to detect delayed inhibitory effects.

Direct antifungal activity against *B. cinerea* was assessed using an automated microbial growth analyser (Bioscreen C MBR^TM^) which monitors optical density in microculture wells over time. Conidia were collected from fresh V8 cultures, filtered, and adjusted to 2 × 10⁵ conidia.mL⁻¹ in 0.6% (w/v) PDB. Each treatment was diluted in the same medium to the desired final concentration. For each condition, 300 µL of conidial suspension containing the treatment were dispensed into three replicate wells of a honeycomb microplate. Border wells were filled with sterile water to avoid edge effects. Plates were incubated at 20°C with continuous shaking inside the analyser. Optical density at 492 nm was recorded every 2 h for approximately 60 h. Growth curves were extracted using the Bioscreen software, and antifungal activity was expressed as the reduction in growth rate or final optical density relative to the untreated control.

### 2.5. ROS measurements

Two leaf discs (3 mm in diameter) were collected from two independent plants and placed in each well of a black 96-well plates (Black 96-Well Immuno Plates, 7605). Three and four technical repeats per treatment were done for grapevine and *A. thaliana*, respectively. Leaf discs were floated overnight in 100 µL of distilled water in the dark.

The following day, the water was replaced by 100 µL of solution containing 200 µM L-012 (Fujifilm, 120-04891) and 200 µg.mL⁻¹ horseradish peroxidase (Sigma-Aldrich, P8125), with or without treatments (Albert et al., 2015). Luminescence was immediately recorded as relative light units (RLU) using a microplate reader (Tecan Infinite M200 PRO), with measurements taken at 2 min intervals and an integration time of 3000 ms over a 60 min period. For NaBut pre-treatment, grapevine leaf discs were incubated in 1 mM NaBut solution for 3 h prior to the addition of the ROS assay solution.

### 2.6. RT-qPCR

For RT-qPCR analyses, treated leaves were collected and ground in liquid nitrogen. Total RNA was isolated using the SV Total RNA Isolation System (Promega), including the DNase treatment step. Reverse transcription was subsequently carried out using the High-Capacity cDNA Reverse Transcription Kit (Applied Biosystems) to generate first-strand cDNA. Quantitative PCR reactions were performed on a ViiA™ 7 Real-Time PCR System (Applied Biosystems) using 10 ng of cDNA and 1× GoTaq^®^ qPCR Master Mix (Promega).

Relative transcript abundance was determined by taking into account the amplification efficiency (E) of each primer pair, as estimated using the LinRegPCR quantitative PCR data analysis software (Ruijter et al., 2009). For *A. thaliana* samples, transcript levels were normalized against two reference genes, *AtRHIP1* (At4g26410) and *AtPTB1* (At3g01150). The sequences of all primers used in this study are provided in Table S2.

### 2.7. Western Blot analysis

For VvHD2 protein immunodetection, 2 g of *A. thaliana* leaves were collected from each genotype. Samples were ground in liquid nitrogen and homogenized in two volumes of nuclear extraction buffer, as described by Dahan et al. (2009). All subsequent steps were performed on ice, and centrifugation steps were carried out at 4°C according to the protocol described by Dahan et al. (2009). The resulting nuclear pellets were resuspended in a high-salt solution containing 500 mM NaCl and 250 mM MgCl_2_ to disrupt the nuclear envelope and release nuclear proteins. For immunoblot analysis, 25 µg of nuclear fraction protein extracts were separated by SDS-PAGE and transferred onto nitrocellulose membranes. Membranes were blocked with 5% milk and incubated with either anti-VvHD2a or anti-VvHD2b polyclonal antibodies. These antibodies were raised in rabbits against recombinant proteins produced in *E. coli* and purified prior to immunization (developed by RD-Biotech, Besançon, France).

For MAPK phosphorylation analysis, leaves from *in vitro*-grown grapevine plantlets were vacuum-preinfiltrated with ultrapure water and left to equilibrate for 2–3 h in six-well plates before treatment. For treatment, the water was replaced with the corresponding treatment solutions. NaBut pre-treatment were done 3 h before treatment. Thirty minutes after treatment, three leaf discs were excised from three independent leaves and immediately frozen in liquid nitrogen, and stored at −80°C. Proteins were extracted from total ground tissue using the TriReagent^®^ protocol described by Marzari et al. (2025). For immunoblot analysis, 20 µg of total protein extracts were separated by SDS-PAGE and transferred onto nitrocellulose membranes. MAPK phosphorylation was detected using an anti-phospho-p44/42 MAPK (Erk1/2) antibody recognizing the Thr202/Tyr204 phosphorylation sites (Cell Signaling Technology, 9101). For both antibodies, immunodetection was performed using ECL Prime Western Blotting Detection Reagent (Cytiva), and chemiluminescence signals were visualized using an Amersham ImageQuant 800 system (Cytiva).

### 2.8. Confocal microscopy

Confocal imaging was performed using a Leica TCS SP8 multiphoton microscope equipped with a 40X oil-immersion objective. Leaf segments from *A. thaliana* lines expressing GFP-tagged VvHD2a or VvHD2b were mounted in ultrapure water between glass slides and coverslips and examined by confocal microscopy. Nuclei were visualized by staining the samples with DAPI prior to imaging. GFP fluorescence was excited at 488 nm, while DAPI fluorescence was excited at 350 nm. GFP and DAPI emissions were detected at 500-525 nm and at 460 nm, respectively. Images were acquired and processed using LAS X software.

### 2.9. Transcriptomic analysis by RNA-seq

For transcriptomic analyses, leaves from *V. vinifera* cv. Marselan plants grown in glasshouses were collected after treatment with NaBut, Belvine^®^, or a combination of both. For each treatment condition, three biological replicates were sampled. The first and second fully expanded leaves were selected, excised at the petiole base, and flash frozen in liquid nitrogen. Samples were stored at −80°C until processing.

Total RNA was extracted from 100 mg of finely ground frozen tissue using the NucleoSpin RNA Plant and Fungi kit (Macherey-Nagel), following the manufacturer’s instructions. RNA quantity and purity were assessed using a Nanodrop spectrophotometer, and RNA integrity was verified using an Agilent Bioanalyzer RNA 6000 Nano chip. Only high-quality RNA samples (RIN > 7) were used for library preparation. High throughput sequencing was performed by BGI (Shenzhen, China); raw reads were filtered and quality-checked prior to delivery. Raw read counts were analysed using the edgeR package in R. Lowly expressed genes were filtered prior to analysis, and library sizes were normalized using the trimmed mean of M-values (TMM) method. Detailed information regarding data collection, validation, and quality control can be found in the accompanying data descriptor (Ilbert et al., associated Data paper).

#### 2.9.1. Differential expression analysis

Differential expression analyses were performed by comparing each treatment (NaBut, Belvine^®^, and NaBut + Belvine^®^) with the water control at 2 h and 4 h after Belvine^®^ treatment. Genes with a Benjamini-Hochberg adjusted p-value (FDR) below 0.05 were considered differentially expressed genes (DEGs). DEGs were functionally classified using MapMan annotations. Custom R scripts were then used to summarize and visualize their mean log_2_ fold changes across MapMan functional subcategories.

#### 2.9.2. Interaction analysis

Interaction analysis was performed using the DiCoExpress pipeline (Lambert et al., 2020) at 2 h and 4 h after Belvine^®^ treatment to investigate whether the transcriptional response to Belvine^®^ was affected by the application of NaBut. Belvine^®^ and NaBut were included in the model as two experimental factors, each defined by two levels: control and treatment. For each factor, the control level comprised all samples that did not receive the corresponding treatment. Differential expression was assessed using a negative binomial generalized linear model, with biological replicates included in the model. Three contrasts were considered: the effect of Belvine^®^ in the absence of NaBut treatment (Bel), the effect of Belvine^®^ in the presence of NaBut treatment (NaBut + Bel), and the interaction between the two treatments (NaBut × Bel), calculated as the difference between these two effects. This interaction contrast identifies genes for which the transcriptional response to Belvine^®^ differed depending on the treatment with NaBut. Genes with a Benjamini-Hochberg adjusted p-value (FDR) below 0.05 were considered to show a significant interaction. Interaction genes were functionally annotated using MapMan, and their functional distribution was visualized using custom R scripts.

### 2.10. Targeted-transcriptomic analysis

Leaves of Cabernet Sauvignon plants from the vineyard experiment conducted in the experimental plot BC2Grape in Bordeaux (44.792498, −0.574980) were collected, frozen in liquid nitrogen and then ground to a powder. Total RNA was extracted using the NucleoMag® RNA Kit (Macherey-Nagel), following the manufacturer’s instructions.

For reverse transcription, 1 µL of oligo(dT) primer was added to 10 µL of RNA per well, followed by incubation at 65°C for 5 min. A volume of 8.5 µL of reverse transcription mix was then added to each well. This mix contained RT buffer, DTT, dNTPs, M-MLV reverse transcriptase, RiboLock RNase inhibitor and RNase-free water. Reverse transcription was carried out at 37°C for 50 min, followed by enzyme inactivation at 70°C for 10 min. The resulting cDNAs were pre-amplified using a multiplex primer pool containing 96 primer pairs, Fluidigm Preamp Master Mix and nuclease-free water according to the manufacturer’s protocol (Standard BioTools^®^). Each reaction contained 3.7 µL of pre-amplification mix and 1.3 µL of cDNA. After initial denaturation, amplification was performed for 20 cycles of 95°C for 15 s and 55°C for 30 s. Residual primers were removed by adding 2 µL of exonuclease I treatment mix to each well. The mix was prepared using nuclease-free water, exonuclease I buffer and exonuclease I. Samples were incubated at 37°C for 30 min, followed by enzyme inactivation at 80°C for 15 min. The resulting products were diluted fivefold in nuclease-free water before real-time quantitative PCR (qPCR) on the BioMark™ HD system (Standard BioTools, formerly Fluidigm), using 96.96 Dynamic Array™ integrated fluidic circuits (IFCs).

Relative gene expression levels were calculated using Pfaffl’s model formula 2^-ΔΔCq^, allowing to consider multiple reference genes for improved normalization.

To compare RNA-seq data from plants grown under controlled conditions with targeted gene expression data from the BC2Grape experimental plot, the two datasets were matched based on common defense-related genes. Genes analyzed using the microfluidic qPCR array in vineyard samples were matched to RNA-seq genes (Vitvi) based on their respective annotations. Vitvi genes were considered as a match when they shared the same functional annotation as the vineyard marker, or as a clearly related member of the same gene family. For multigenic families a single vineyard marker could match multiple Vitvi genes. The complete list of marker-Vitvi associations is provided in Table S3.

At each RNA-seq time point (2 and 4 h), only Vitvi genes significantly differentially expressed (FDR < 0.05) in both Belvine^®^ and NaBut + Belvine^®^ conditions were retained. The effect of NaBut was calculated as log2FC(NaBut + Belvine^®^) - log2FC(Belvine^®^). In the vineyard experiment, the effect of NaBut was calculated at each sampling time by subtracting the mean expression (Copper + Belvine^®^) to the mean expression (Copper + Belvine^®^ + NaBut).

For each matched marker-Vitvi pair, the NaBut effect at each RNA-seq time point was compared with the NaBut effect at each vineyard sampling time. Each of these RNA-seq– vineyard comparisons were included in the concordance analysis. A comparison was considered concordant when NaBut had the same effect in both experiments (positive or negative), and discordant when they had opposite effects. Because sampling kinetics differed between the two experiments, RNA-seq and vineyard time points were not considered direct temporal equivalents.

### 2.11. Vineyard protections experiments

Field experiments were conducted on Chardonnay grapevines at two vineyard sites in Burgundy (France): Marsannay-la-Côte (Côte-d’Or, 47.2622, 4.9900) and Azé (Saône-et-Loire, 46.4232, 4.7553). At both sites, trials followed a randomized block design with three blocks per treatment. Five treatments were compared: a water-treated control, copper hydroxide (Champ® Flo Ampli, Nufarm) with a full seasonal copper input of 3.9 kg Cu/ha, a reduced-copper program, a reduced copper combined with Belvine^®^, and the same combination supplemented with NaBut (220 mg/L). The reduced-copper program differed between sites. At Marsannay-la-Côte, copper applications were reduced by 50% relative to the full seasonal copper program throughout the growing season. At Azé, the 50% copper reduction was applied before flowering, while a conventional copper-based program using Bordeaux mixture was applied from flowering onwards. Belvine^®^ application rates also differed between sites, with 0.75 g/L applied at Marsannay-la-Côte and 1.625 g/L at Azé, the latter corresponding to half the standard field application rate (3.25 g/L).

At Marsannay-la-Côte, a combined “field-laboratory” approach was used. Treatments were applied to grapevines grown in the vineyard, after which leaves were collected 48 h later for *P. viticola* inoculation in the laboratory. Three independent experiments were performed over several weeks, each involving a separate leaf sampling and inoculation. Leaf discs were excised and placed on moistened Whatman paper. After 2 h, the discs were inoculated with a *P. viticola* suspension (1 × 10⁵ sporangia mL⁻¹), sprayed onto the abaxial leaf surface to produce a thin, homogeneous film of droplets (using a Cosmos 18 Pro spayer, Berthoud, France). Following inoculation, leaf discs were maintained in darkness at 100% relative humidity for at least 6 h to promote zoospore release and infection, and subsequently incubated under a 10 h/14 h light/dark cycle at 20/18°C and 100% relative humidity to induce sporulation. Disease severity was assessed at 6, 7 and 8 dpi by imaging the leaf discs and quantifying the sporulating area using Visilog 6.9 image analysis software (Kim Khiook et al., 2013). At Azé, the strategy was evaluated entirely under vineyard conditions over two consecutive growing seasons. Treatments were applied by the grower, and *P. viticola* infection was promoted by artificial inoculation combined with misting. DM severity on leaves and bunches was visually scored as the percentage of diseased tissue per leaf or bunch on a 0-100% scale. Assessments were performed at several time points during each growing season by scoring 100 leaves and 50 bunches per block at each assessment date. For statistical analysis, one assessment date was selected for each organ and year, corresponding to a time point when disease severity was sufficiently high to allow treatment efficacy to be evaluated, while avoiding complete disease saturation across all treatments.

Disease severity was analysed within a Bayesian framework using generalized regression models implemented in the brms package in R. A zero-one-inflated Beta distribution was used to account for proportional data including observations at 0 and 1. At Marsannay-la-Côte, leaf severity was modelled as Severity ∼ Treatment + (1 | Block). At Azé, leaf severity was modelled across both growing seasons as Severity ∼ Treatment + Year + (1 | Block), with year included as a fixed effect and block as a random effect. Bunch severity was modelled as Severity ∼ Treatment + Year, without a random block effect because separate block-level observations were not available for bunches during the first growing season. Two complementary analyses were derived from these models. First, the specific effect of NaBut supplementation was assessed by comparing reduced copper + Bel + NaBut with reduced copper + Bel, and was summarized by the posterior probability of improved protection. Second, model-adjusted severity estimates were generated for each treatment to compare the performance of the reduced copper + Bel + NaBut program with the full-rate and reduced-copper references. For Azé, estimates were averaged with equal weight across the two growing seasons to provide interannual estimates. Weakly informative Normal (0, 1.5) priors were specified for regression coefficients, and Exponential (1) priors for random effect standard deviations where applicable. Model convergence and fit were assessed using R^, bulk and tail effective sample sizes, divergent transitions, maximum tree depth, and posterior predictive checks.

## 3. Results

### 3.1. Grapevine genome contains and expresses two HD2 orthologues

Bourque et al. (2016) identified two HD2 orthologues in grapevine. With the availability of updated grapevine genome assemblies, we sought to determine whether additional HD2 family members were present in this species. To this end, we performed a BLAST protein search and still identified only two proteins, namely VvHD2a and VvHD2b. We constructed a phylogenetic tree including all HDACs identified in *A. thaliana*, subdivided into the three major subfamilies (RPD3/HDA1, SIR2, and HD2), as well as HD2 members from *Oryza sativa*, *Nicotiana tabacum*, *Vitis riparia* and *V. vinifera*. As expected, our analysis showed that VvHD2a/b segregates with other HD2s (Fig. 1A, Table S4). From a structural perspective, both grapevine HD2 proteins contain the conserved domains previously described by Bourque et al. (2016), including a conserved N-terminal HD2 specific-label, a catalytic domain in N-Ter, a central acidic regulatory domain (containing conserved sites specific to MAPKs and Casein Kinase 2α), a nuclear localization signal (NLS), and a TFIIIA-type zinc finger in C-Ter (Fig. 1B, S1). We next investigated the functional characteristics of these two HD2s. First, VvHD2a and VvHD2b were constitutively expressed with GFP fusion proteins in *A. thaliana* to confirm the nuclear localization predicted from the presence of the conserved NLS motif. For both proteins, GFP fluorescence co-localized with DAPI staining, confirming their nuclear localization. Notably, a stronger fluorescence signal was observed within the nucleolus, indicating that both proteins are correctly targeted to their expected subcellular compartment in the heterologous *A. thaliana* system (Figure 1C-D).

**Figure 1.**
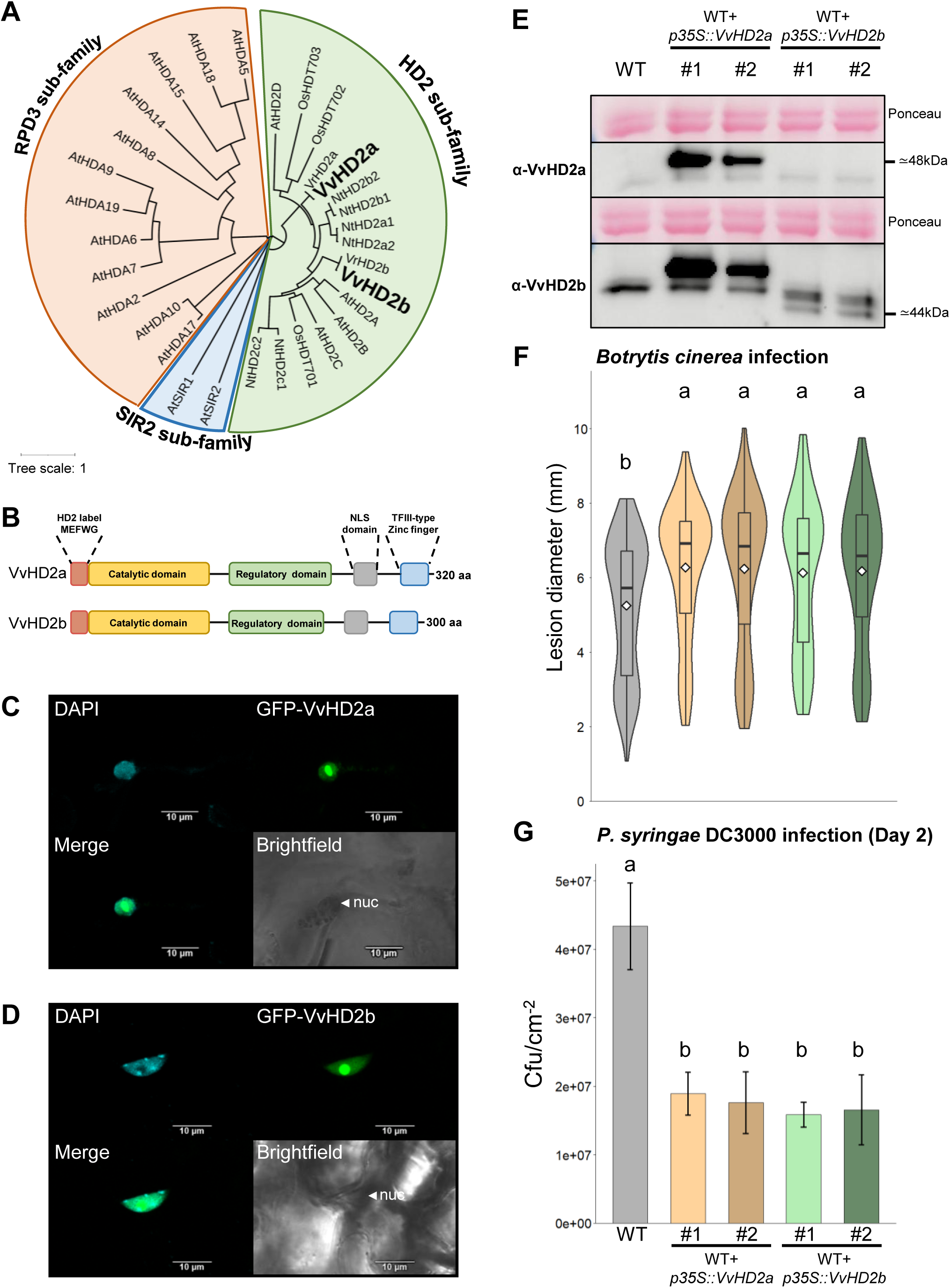
Histone deacetylases of type 2 (HD2) from *Vitis vinifera* are involved in plant immunity. (A) Maximum-likelihood phylogenetic tree constructed using the Jones Taylor Thornton (JTT) substitution model with 1000 bootstrap replicates. The tree includes the 18 histone deacetylases (HDACs) from *Arabidopsis thaliana* (At), grouped into the RPD3 (red), SIR2 (blue), and HD2 (green) subfamilies, together with HD2 proteins from *Oryza sativa* (Os), *Nicotiana tabacum* (Nt), and *Vitis vinifera* (Vv). The phylogenetic analysis was performed using MEGA XII (Kumar *et al*., 2024). (B) Schematic representation of the predicted domain organization of VvHD2a and VvHD2b. The red box indicates the highly conserved N-terminal domain specific to the HD2 subfamily; the yellow box corresponds to the catalytic domain containing highly conserved regions; the green box represents the regulatory domain harboring putative CK2a and MAPK phosphorylation sites; the grey box indicates a KKxK monopartite nuclear localization signal (NLS); and the blue box corresponds to the C2H2-type zinc-finger motif described by Bourque *et al*. (2016). (C, D) Subcellular localization of GFP-tagged VvHD2a (C) and VvHD2b (D) in *A. thaliana* leaf epidermal cells. GFP fluorescence co-localized with DAPI staining, indicating nuclear localization. Scale bars = 10 µm. Similar localization patterns were observed in two independent transgenic lines for each construct. (E) Accumulation of VvHD2a and VvHD2b proteins in WT plants and two independent transgenic lines constitutively expressing either VvHD2a (WT+*p35S::VvHD2a#1,#2)* or VvHD2b (WT+*p35S::VvHD2b#1,#2)*, as determined by immunoblot analysis using polyclonal antibodies raised against the corresponding recombinant proteins. Twenty micrograms of nuclear proteins, extracted as described in the Materials and Methods section, were loaded per lane. (F) Lesion diameter induced by *Botrytis cinerea* on WT plants (grey), WT+*p35S::VvHD2a#1,#2* (light and dark orange) and WT+*p35S::VvHD2b#1,#2* (light and dark green), previously validated by immunoblot analysis, measured 3 days post-inoculation (dpi). Violin plots with embedded boxplots show the distribution of lesion diameters from four independent biological experiments. For each biological replicate, 30 lesions were scored from nine plants per genotype. (G) Bacterial growth following infection with *Pseudomonas syringae* pv. *tomato* DC3000. Bars represent the mean (± SE) colony-forming units (CFU) per cm² of leaf tissue in WT plants (grey), WT+*p35S::VvHD2a#1,#2* (light and dark orange) and WT+*p35S::VvHD2b#1,#2* (light and dark green), measured 2 dpi (Day 2) from three independent biological experiments. For (F) and (G) different letters indicate statistically significant differences among genotypes (Kruskal–Wallis test followed by Dunn’s multiple comparison test with Bonferroni correction, *P* < 0.05).

### 3.2. HD2s from *Vitis vinifera* are involved in the control of plant immunity

To further investigate the roles of VvHD2a and VvHD2b, we constitutively overexpressed each gene, without a GFP tag, in the *A. thaliana* Col-0 ecotype rather than in an HD2 mutant background. This strategy was adopted because *A. thaliana* possesses four HD2 genes, making the generation of a quadruple mutant technically challenging. To verify the accumulation of the transgenic proteins in the selected homozygous lines, we used polyclonal antibodies raised against recombinant VvHD2a and VvHD2b proteins. As shown in Fig. 1E, the α-VvHD2a antibody specifically recognized VvHD2a and did not detect any signal in wild-type (WT) plants nor in lines expressing VvHD2b. In contrast, the α-VvHD2b antibody recognized both VvHD2 proteins and potentially endogenous HD2 proteins from *A. thaliana*, as indicated by the presence of a signal in WT plants. This cross-reactivity is likely due to conserved epitopes shared between these closely related proteins (69.6% of similarity between VvHD2a and VvHD2b, Fig. 1E, S1). Furthermore, the α-VvHD2b antibody confirmed the predicted size difference between VvHD2a and VvHD2b (Fig. 1B), based on their apparent molecular masses, estimated at approximately 48 and 44 kDa, respectively. Immunoblot analyses clearly confirmed the accumulation of the transgenic proteins in the corresponding transgenic lines, demonstrating that the transgenes were stably expressed and not subjected to silencing. Following confirmation of transgene expression, we investigated the involvement of VvHD2a and VvHD2b in plant immunity using two distinct pathogens. First, we assessed the response of the transgenic lines to infection by the necrotrophic fungus *B. cinerea*. The mean lesion diameter was 5.25 mm in WT plants, whereas it ranged from 6.13 to 6.26 mm across all transgenic lines. This phenotype was consistently observed across independent transgenic lines expressing either of the two grapevine HD2 proteins, indicating that overexpression of VvHD2a or VvHD2b enhances significantly plant susceptibility to this necrotrophic pathogen compared with WT plants (Fig. 1F). We next investigated the impact of constitutive VvHD2 expression on the immune response to *Pseudomonas syringae* pv. *tomato*, a pathogen that infects plants through a different mode of infection. Inoculation was done at about 3.0 × 10^3^ CFU.cm⁻² of leaf area (Fig. S2). Unexpectedly, after two days, transgenic lines overexpressing VvHD2a or VvHD2b exhibited enhanced resistance to this bacterial pathogen, with bacterial populations remaining below 2.0 × 10⁷ CFU.cm⁻² in all transgenic lines, compared with 4.3 × 10⁷ CFU.cm⁻² in WT plants (Fig. 1G). These results indicate that the contribution of VvHD2 proteins to plant immunity depends on the mode of action of the pathogen, suggesting that they can act as either positive or negative regulators of plant immune responses depending on the pathosystem.

### 3.3. Constitutive expression of VvHD2 proteins reduces flg22 triggered immune responses in *A. thaliana*

To further elucidate the functions of VvHD2a and VvHD2b, we investigated early PTI responses induced by flg22 by monitoring ROS burst. Transgenic lines overexpressing *VvHD2a* or *VvHD2b* displayed a significant reduction in the amplitude of the flg22-induced ROS burst compared with WT plants, which reached a peak of approximately 10,500 RLU (Fig. 2A, D). In the two *VvHD2a*-overexpressing lines, ROS production peaked at approximately 5,000 RLU, whereas the two *VvHD2b*-overexpressing lines reached maximum levels of approximately 4,300 and 6,300 RLU, respectively (Fig. 2A, D). Consistent with these observations, quantification of the total area under the ROS production curve revealed a significant reduction in ROS accumulation in all four transgenic lines compared with WT plants (Fig. 2B, E). Overall, these results indicate that constitutive expression of either *VvHD2a* or *VvHD2b* attenuates the flg22-induced ROS burst in *A. thaliana*, suggesting that both proteins act as negative regulators of this early PTI response.

**Figure 2.**
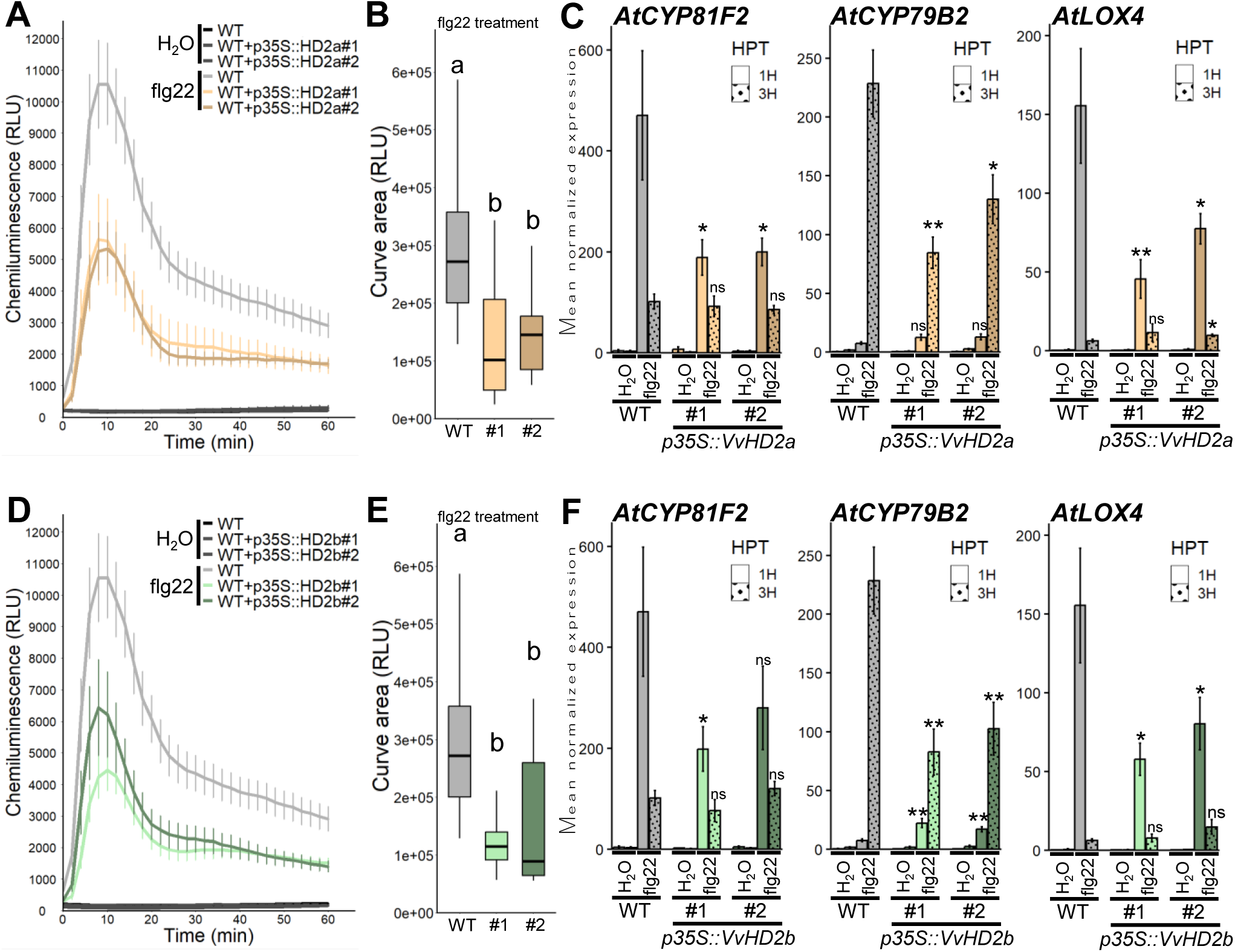
VvHD2a and VvHD2b are involved in PTI response in *A. thaliana*. (A-B, D-E) ROS burst detected by the luminol-horseradish peroxidase approach in WT (grey) and transgenic lines either overexpressing *VvHD2a* (light and dark orange) or *VvHD2b* (light and dark green). (A) ROS burst of WT and WT+*p35S::VvHD2a#1,#2* transgenic lines treated with H_2_O or flg22. (B) Box plots representing curve area corresponding to total ROS accumulation during 1 hour comparing WT and WT+*p35S::VvHD2a#1,#2* following flg22 treatment. (D) ROS burst of WT and WT+*p35S::VvHD2b#1,#2* transgenic lines treated with H_2_O or flg22. (E) Box plots representing curve area corresponding to total ROS accumulation during 1 hour comparing WT and WT+*p35S::VvHD2b#1,#2* following flg22 treatment. RLU: relative luminescence unit. For (B) and (E) different letters indicate statistically significant differences among genotypes (Kruskal–Wallis test followed by Dunn’s multiple comparison test with Bonferroni correction, *P* < 0.05). (C, F) Normalized expression level of *AtCYP81F2, AtCYP79B2* and *AtLOX4* measured by qPCR 1h and 3h after flg22 (1µM) or H_2_O treatment. The expression levels of *AtCYP81F2, AtCYP79B2* and *AtLOX4* were normalized to those of two housekeeping genes. Bars represent the mean of relative expression ± SEM of 6 biologically independent experiments. Asterisks indicate a statistically significant difference between each transgenic line and the WT treated with flg22, (non-parametric Wilcoxon test; *, P<0.05; **, P<0.01). (C) Normalized expression level of *AtCYP81F2, AtCYP79B2* and *AtLOX4* comparing normalized expression level between WT and WT+*p35S::VvHD2a#1,#2* transgenic lines (light and dark orange). (F) Normalized expression level of *AtCYP81F2, AtCYP79B2* and *AtLOX4* comparing normalized expression level between WT and WT+*p35S::VvHD2b#1,#2* transgenic lines (light and dark green).

In addition, we assessed the expression of three defense-related genes induced by flg22 treatment: *AtCYP81F2* (*CYTOCHROME P450 FAMILY 81 SUBFAMILY F POLYPEPTIDE 2*) and *AtCYP79B2* (*CYTOCHROME P450 FAMILY 79 SUBFAMILY B POLYPEPTIDE 2*), two genes involved in indole glucosinolate biosynthesis (Denoux et al., 2008), and *AtLOX4* (*LIPOXYGENASE 4*), a gene involved in jasmonic acid biosynthesis (Denoux et al., 2008; Yang et al., 2020). In transgenic lines constitutively expressing *VvHD2a*, flg22-induced expression of *AtCYP81F2* and *AtLOX4* was significantly reduced by more than 50% in both independent lines 1 h after treatment. Likewise, *AtCYP79B2* transcript accumulation was significantly reduced by approximately 40-60%, depending on the transgenic line, 3 h after flg22 treatment (Fig. 2C). A similar expression pattern was observed in *VvHD2b* overexpressing lines. The induction of *AtCYP81F2* and *AtLOX4* was reduced by more than 50% 1 h after flg22 treatment in both lines. However, although *AtCYP81F2* expression tended to be lower in line 2, this reduction was not statistically significant. In addition, both *VvHD2b* overexpressing lines exhibited a significant reduction (>50%) in *AtCYP79B2* expression 3 h after flg22 treatment (Fig. 2F).

Together, these results indicate that constitutive expression of *VvHD2a* and *VvHD2b* attenuates the transcript abundance of several flg22-responsive defense genes, further supporting their role as negative regulators of early pattern-triggered immune responses.

### 3.4. NaBut reverses the increased susceptibility to *B. cinerea* induced by *VvHD2a* or ***VvHD2b* overexpression.**

Our results consistently demonstrate that VvHD2a and VvHD2b act as negative regulators of plant immunity, particularly during PTI. In mammals, NaBut is a well-characterized HDAC inhibitor (Steliou et al., 2012). We therefore sought to determine whether NaBut could counteract the effects of constitutive *VvHD2a* and *VvHD2b* expression. To this end, we assessed the effects of NaBut treatment on WT *A. thaliana* plants and transgenic lines overexpressing either *VvHD2a* or *VvHD2b*. Plants were treated with NaBut prior to inoculation with *B. cinerea*. NaBut treatment did not significantly affect lesion development in WT plants (Fig. 3). In contrast, NaBut significantly reduced lesion size in both *VvHD2a*- and *VvHD2b*-overexpressing lines. In *VvHD2a*-overexpressing lines, lesion diameters decreased from approximately 5.4 mm in water-treated plants to 4.0 mm in NaBut-treated plants. Notably, NaBut-treated *VvHD2a*-overexpressing lines exhibited significantly smaller lesions than NaBut-treated WT plants (Fig. 3). In *VvHD2b*-overexpressing lines, lesion diameters decreased from 5.5-5.6 mm in water-treated plants to approximately 4.3 mm following NaBut treatment (Fig. 3). However, unlike *VvHD2a*-overexpressing lines, NaBut-treated *VvHD2b*-overexpressing lines did not differ significantly from NaBut-treated WT plants in terms of lesion size.

**Figure 3.**
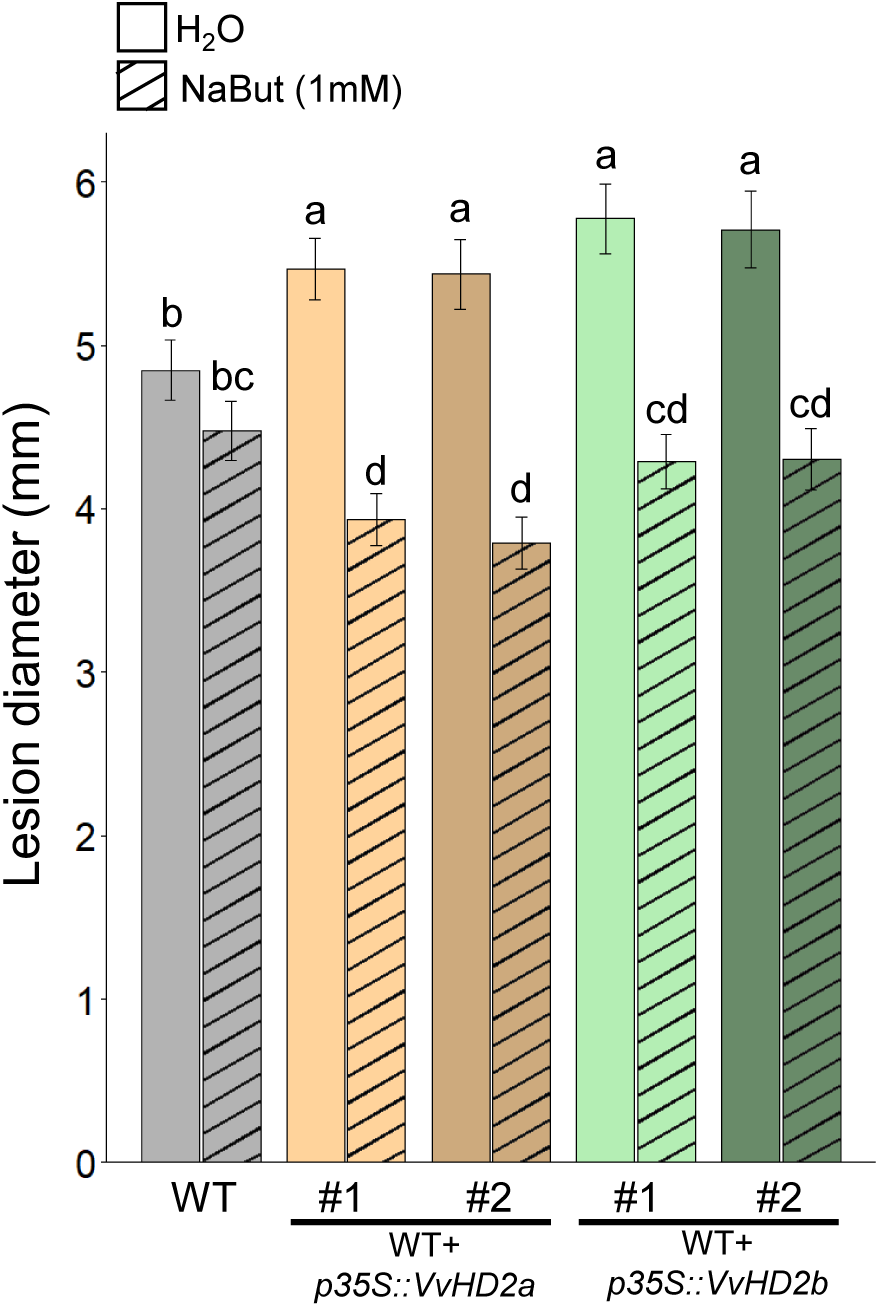
Sodium butyrate (NaBut) treatment reverses the enhanced susceptibility of *VvHD2a*- and *VvHD2b*-overexpressing Arabidopsis lines to *Botrytis cinerea*. Bar plots represent lesion diameter induced by *Botrytis cinerea* on WT plants (grey), WT+*p35S::VvHD2a#1,#2 (*light and dark orange) and WT+*p35S::VvHD2b#1,#2* (light and dark green), treated with H_2_O or NaBut (1mM) 24h before *B. cinerea* inoculation and measured 3 dpi. Different letters indicate statistically significant differences among genotypes (Kruskal–Wallis test followed by Dunn’s multiple comparison test with Benjamini–Hochberg correction, *P* < 0.05).

These results indicate that NaBut suppresses the enhanced susceptibility associated with constitutive expression of *VvHD2a* and *VvHD2b* during *B. cinerea* infection. Consequently, these findings support the potential use of HDAC inhibitors such as NaBut as a strategy to block or reduce VvHD2a- and VvHD2b-mediated repression of plant immunity and ultimately enhance disease resistance.

### 3.5. In grapevine NaBut improves resistance to *B. cinerea* and enhances PDI-induced protection against *P. viticola*

NaBut treatment significantly enhanced grapevine (cv. Marselan) resistance to *B. cinerea*, reducing lesion area by 37 % compared with the water-treated control (Fig. 4A). This is consistent with the enhanced resistance observed in *Arabidopsis* transgenic lines overexpressing *VvHD2a* and *VvHD2b* following NaBut treatment (Fig. 3). To assess the effect of NaBut against another major grapevine pathogen, we investigated the protection induced against *P. viticola*. Contrary to its protective effect against *B. cinerea*, NaBut did not enhance grapevine resistance to *P. viticola*. Instead, NaBut-treated plants exhibited a tendency toward increased disease development compared with the H_2_O-treated control (average protection index of −35.6 %), although this difference was not statistically significant (Fig. 4B). Previous work by Bourque et al. (2011) showed that combining NaBut with a pathogen-derived elicitor, such as the oomycete elicitor cryptogein, enhanced defense activation compared with the elicitor treatment alone. Based on these findings, we investigated whether NaBut could similarly enhance elicitor-induced protection against *P. viticola*. To this end, we used the commercial biocontrol product Belvine^®^, which is based on a yeast extract and is approved for use against *P. viticola*. Grapevine plants of *V. vinifera* cv. Marselan were treated with NaBut and Belvine^®^, either alone or in combination, 48 h before pathogen inoculation. Whereas Belvine^®^ applied at 3.25 g.L^-1^, corresponding to the recommended field rate, provided a high level of protection, with a protection index of 69.5%, Belvine^®^ applied alone at a four-fold lower rate (0.75 g.L^-1^) provided limited protection (9.6%). Remarkably, combining NaBut (1 mM) with a reduced dose of Belvine^®^ (0.75 g.L^-1^) resulted in a high level of protection (66%) similar to that observed with the high dose of Belvine^®^ (Fig. 4B, left panel). To determine whether the protective effect of the combined treatment exceeded that expected from the individual treatments, we compared the observed protection with the theoretical additive effect calculated according to the Colby model (Colby, 1967). The protection observed for the NaBut + Belvine^®^ combination was significantly higher than the expected additive protection (66.0% versus 26.9%, respectively; *P* = 0.044; Fig. 4B, right panel), demonstrating a synergistic interaction between NaBut and Belvine^®^ in the protection of grapevine against *P. viticola*.

**Figure 4.**
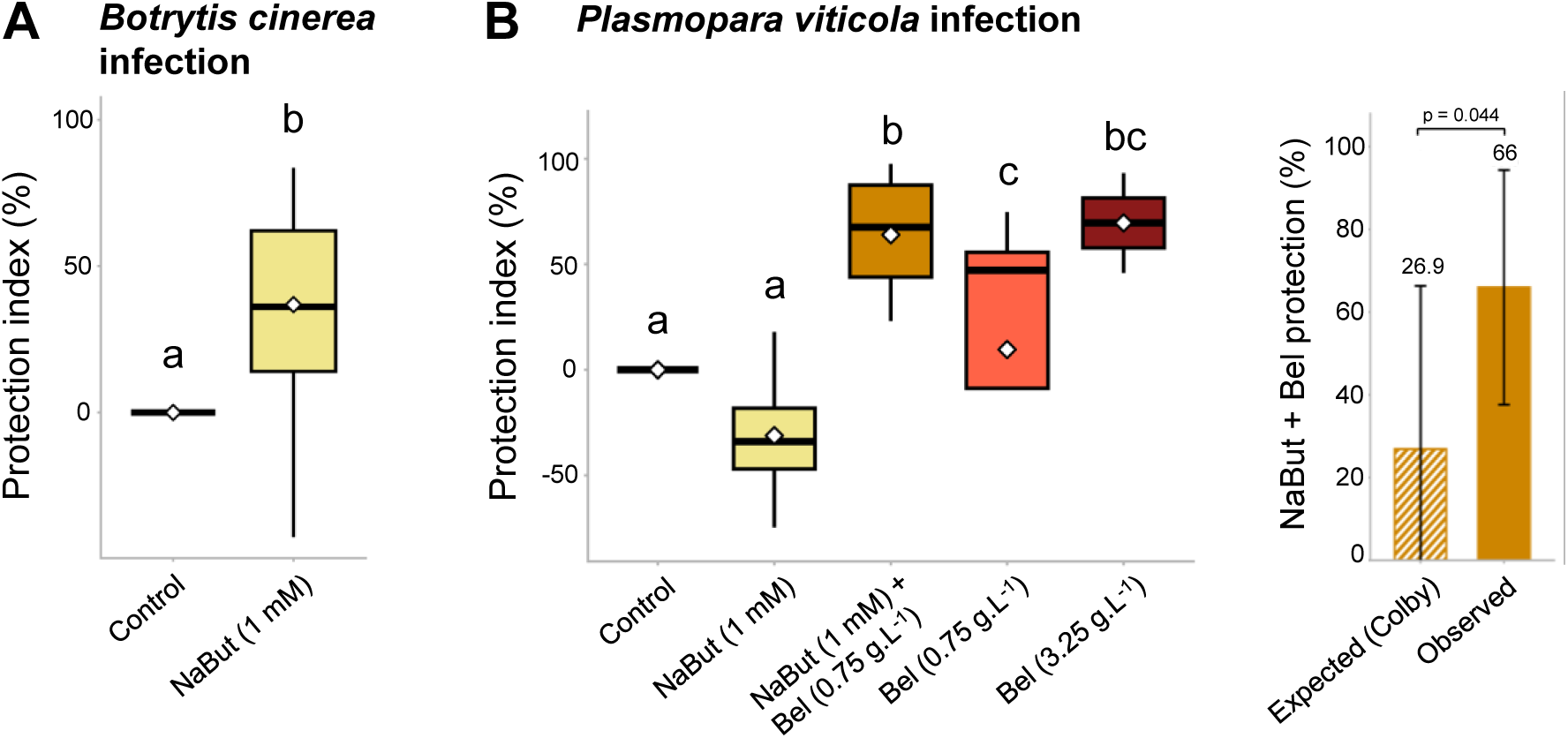
NaBut alone and in combination with Belvine^®^ (PDI) enhance the resistance of grapevine to *Botrytis cinerea* and *Plasmopara viticola*. **(A)** Development of *B. cinerea* at 3 dpi on grapevine leaf discs treated 48 h before with NaBut (1 mM) and H_2_O (Control). Necrosis area were measured 3 dpi and results are expressed as a protection index relative to the H_2_O- treated control. (**B**) Protection against *P. viticola* was assessed by measuring sporulation 5 dpi on leaf discs and expressed as a protection index relative to the H_2_O-treated control (left panel). Grapevine herbaceous cuttings were treated with NaBut (1 mM), Belvine^®^ (0.75 g.L^-1^ and 3.25 g.L^-1^), or H_2_O (Control). For the combination treatment, NaBut was applied 3 h before Belvine^®^ (0.75 g.L^-1^). All treatments were applied 48 h before pathogen inoculation. For both pathogen assays different letters indicate significant differences among treatments (Kruskal–Wallis test followed by Dunn’s multiple-comparison test with Holm’s correction, P < 0.05). The right panel compares the protection expected according to the Colby additivity model with the observed protection obtained for the NaBut Belvine^®^ combination. Statistical significance was assessed using a one-sided linear mixed-effects model. Bars represent the mean ± SD of four independent biological experiments.

To further investigate the mechanisms underlying the protection observed *in planta*, we assessed whether NaBut and Belvine^®^ could directly affect the pathogen motility or growth under *in vitro* conditions. NaBut caused a significant but limited reduction in *B. cinerea* growth at concentrations ranging from 1 to 10 mM (approximately 20% to 35%, depending on the concentration). Regarding *P. viticola*, zoospore motility was reduced only at the highest NaBut concentration tested (10 mM), whereas no significant effect was observed at 1 mM (Fig. S3A-B). Belvine^®^, whether applied alone or in combination with NaBut (1 mM), almost completely inhibited *P. viticola* zoospore motility at both 3.25 and 0.75 g·L⁻¹ (Fig. S3C). This observation is consistent with previous EFSA assessments (European Food Safety Authority (EFSA) et al., 2018). Notably, despite this strong direct inhibitory effect at 0.75 g·L⁻¹, Belvine^®^ provided only limited protection when applied alone *in planta* (9.6%). One has to mention that in protection experiments *P. viticola* was inoculated 48 h after Belvine^®^ treatment; this delay is supposed to allow PDI components to diffuse across the cuticle and reach their target cells. Although a contribution of direct pathogen inhibition to the protective effect of Belvine^®^ cannot be excluded, the strong inhibitory effect of Belvine^®^ alone made it difficult to assess whether NaBut could further enhance this direct activity. These findings suggest that additional mechanisms may contribute to the enhanced protection observed *in planta*. To further investigate these mechanisms, we next examined several plant defense responses.

### 3.6. NaBut selectively potentiates Belvine^®^-induced MAPK phosphorylation without enhancing the ROS burst in grapevine

To investigate whether the NaBut-Belvine^®^ combination also enhances grapevine-mediated immune signaling, we examined two early defense signaling events: the oxidative burst and MAPK activation. NaBut alone did not induced ROS production. Conversely, Belvine^®^ alone triggered a transient ROS burst, reaching a maximum of approximately 700 RLU at 20 min (Fig. 5A). Pretreatment with NaBut for 3 h before Belvine^®^ application did not further increase the amplitude of this response, which remained comparable to that observed with Belvine^®^ alone. Consistent with these results, the area under the curve during the first hour was not significantly different between the two treatments (Fig. 5B). We next analyzed MAPK phosphorylation as a second early signaling event. We did not detect any MAPK phosphorylation in the control nor in NaBut-alone treatments. In contrast, Belvine^®^ induced a slight phosphorylation of two MAPKs after 30 min (45 and 49 kDa). Notably, NaBut pretreatment, before Belvine^®^ treatment induced a stronger phosphorylation of both MAPKs than Belvine^®^ alone (Fig. 5C).

**Figure 5.**
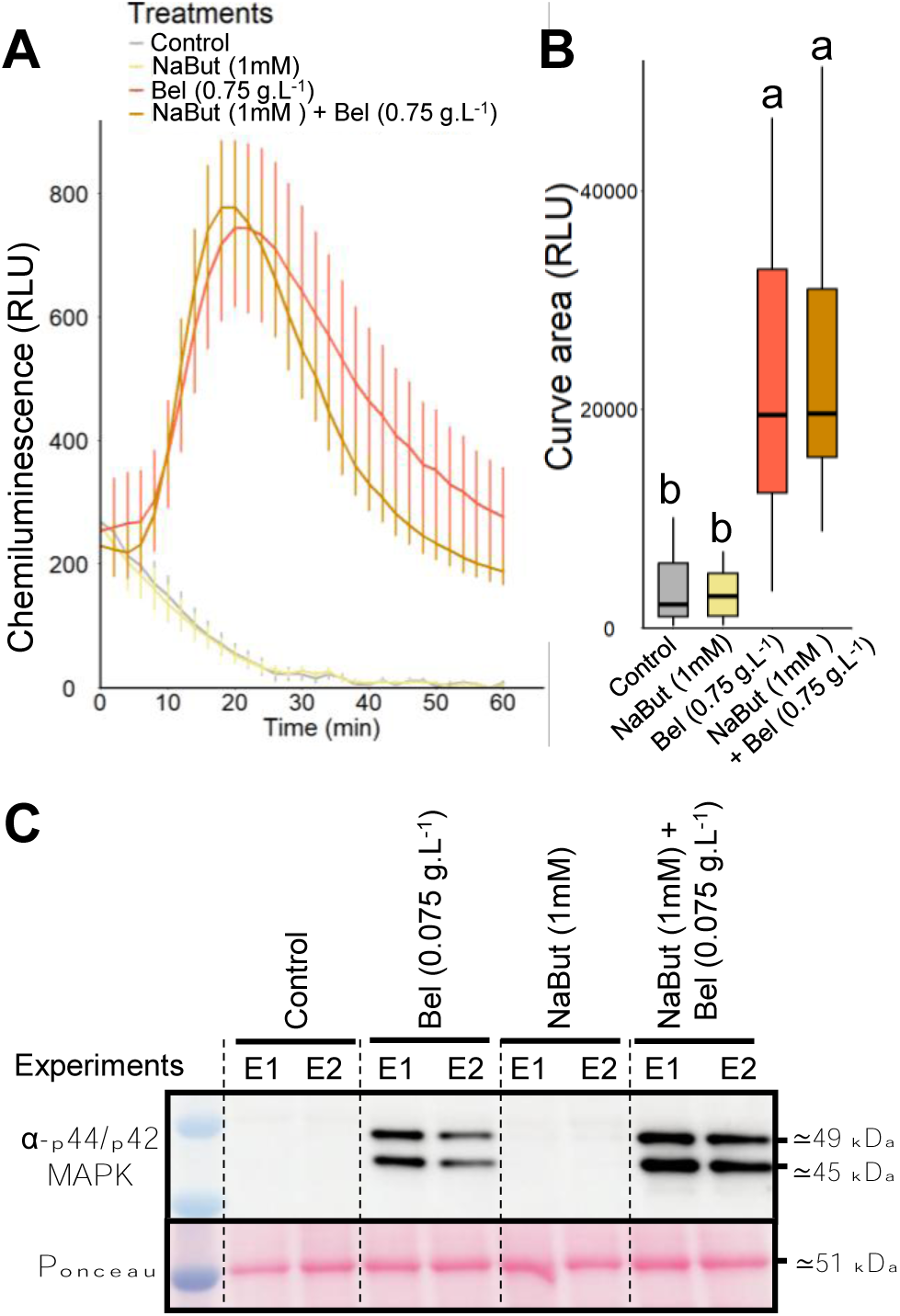
In grapevine, combination of NaBut and Belvine^®^ enhances specific early defense signaling events compared with individual treatments. **(A)** Reactive oxygen species (ROS) burst measured using the luminol–horseradish peroxidase assay in grapevine leaves treated with H_2_O (control, gray), sodium butyrate (NaBut, yellow), Belvine^®^ (Bel, red), or the combination of NaBut and Belvine ^®^ (orange). **(B)** Box plots showing the area under the ROS production curve, corresponding to total ROS accumulation over 1 h, in grapevine leaves treated with H_2_O (control, gray), NaBut (yellow), Bel (red), or the combination of NaBut and Bel (brown). For each experiment, leaf samples were collected from two independent plants. Data represent the mean of eight independent biological experiments. **(C)** Representative immunoblots from two independent experiments (#1 and #2) showing MAPK phosphorylation 30 min after treatment with H_2_O (control), Belvine ^®^ (Bel), NaBut, or the combination of NaBut and Belvine^®^. Phosphorylated MAPKs were detected using an antibody raised against the phosphorylated form of human extracellular signal-regulated kinase 1/2 (α-pERK1/2). For each experiment, leaf samples were collected from three independent plantlets. Similar results were obtained in three independent biological replicates.

Together, these results show that NaBut affects the early immune signaling triggered by Belvine^®^ by specifically enhancing MAPK phosphorylation, but not the ROS burst amplitude.

### 3.7. Transcriptomic analysis reveals that NaBut modulates the Belvine^®^-induced defense response

To investigate the transcriptional changes associated with the synergistic effect observed between NaBut and Belvine^®^, RNA-seq analyses were performed to characterize the responses induced by NaBut and Belvine^®^, applied individually or in combination, particularly in defense-related functional categories, and to determine whether the treatment with NaBut altered the transcriptional response to Belvine^®^.

A global and exploratory overview of MapMan functional subcategories was generated based on DEGs identified from comparisons between single or combined treatments and their respective water controls at 2 h and 4 h (Fig. 6A, Table S5). Overall, both NaBut and Belvine^®^ treatments induced differential gene expression relative to the water control, with patterns varying according to the subcategories. NaBut treatment was more frequently associated with negative mean log_2_ fold changes whereas Belvine^®^ more often induced positive regulation. Under the combined treatment, transcriptional changes generally followed the direction of regulation observed with Belvine^®^, with increased magnitude and/or persistence compared to Belvine^®^ alone, whereas this pattern was not observed with NaBut alone. This overall transcriptional pattern was consistent with the synergistic protective effect observed *in planta*, suggesting that this effect is also reflected at the transcriptional level. Interestingly, this tendency was observed for functional groups linked to immunity, including pathogen response within external stimulus response, reactive oxygen generation within redox homeostasis, ethylene and jasmonic acid within phytohormone action, or lignin metabolism within cell wall organization. Notably, a few subcategories displayed down regulation under NaBut alone but stronger up regulation under the combined treatment compared to Belvine^®^ alone, as observed for ethylene and lignin metabolism (Fig. 6A). Together, these observations suggest that Belvine^®^ may drive the direction of the transcriptional response, while NaBut appears to modulate its magnitude.

**Figure 6.**
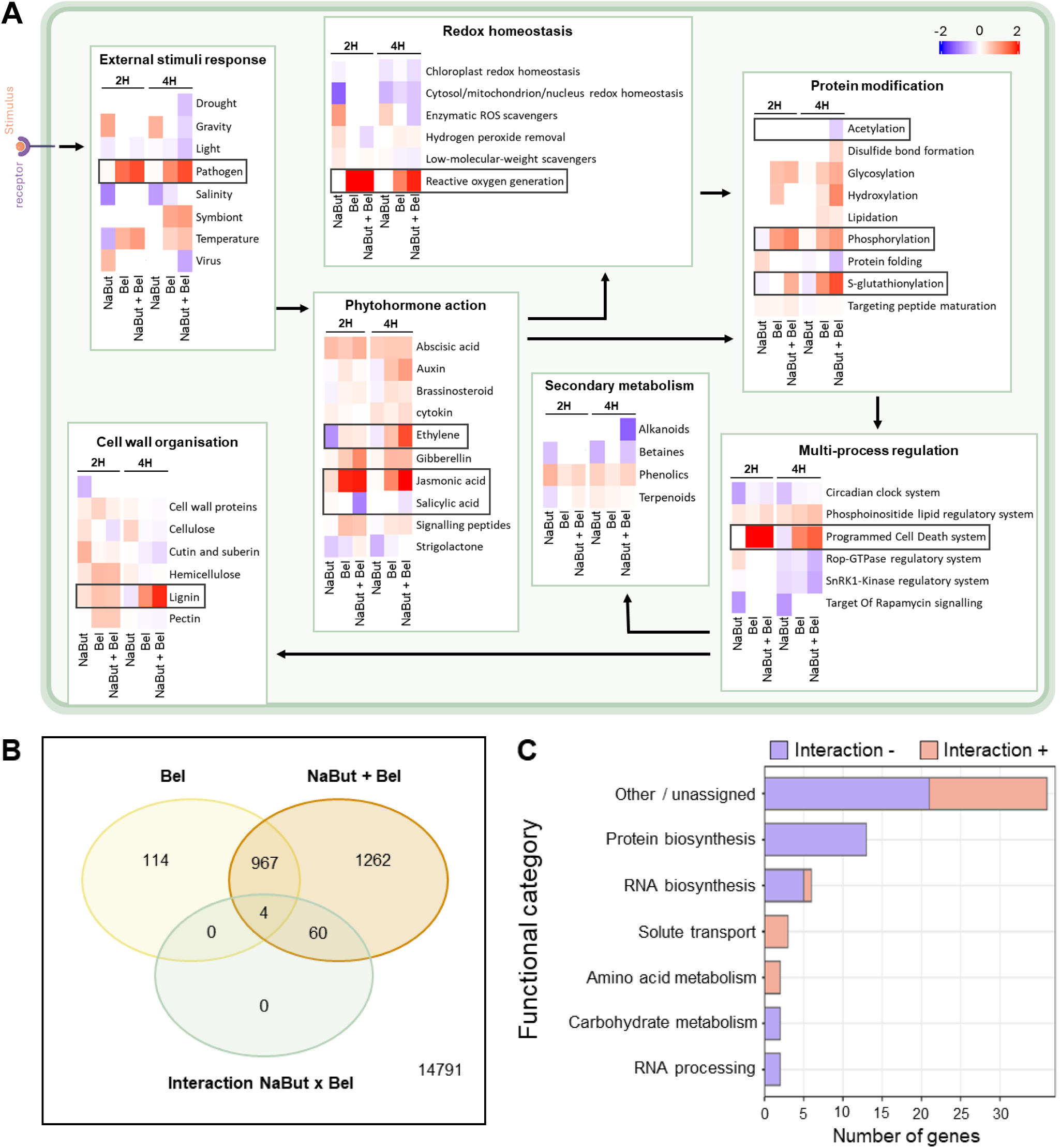
NaBut modulates the grapevine transcriptional response to Belvine^®^. (**A)** Transcriptomic analysis of grapevine leaves following treatment with sodium butyrate (NaBut) and Belvine**^®^** (Bel) alone or a double treatment (NaBut + Bel) consisting of a 3h pre-treatment with NaBut followed by Bel, compared with their corresponding H_2_O controls. Samples were collected 2h and 4h after Bel application. All conditions were harvested simultaneously to allow direct comparison of transcriptional responses. Heatmaps display the mean log_2_ fold change of differentially expressed genes (DEGs : FDR < 0.05) aggregated by MapMan functional categories. Color intensity reflects the magnitude of transcriptional regulation (blue: down-regulation, red: up-regulation; fixed scale from −2 to +2). Dark boxes highlight defense-related functional categories discussed in the text. **(B)** Venn diagram showing the number of DEGs identified at 2 h in Belvine® alone (Bel), Belvine^®^ effect in the presence of NaBut (NaBut + Bel), and genes displaying a significant NaBut × Belvine^®^ interaction effect (NaBut + Bel). Differential expression analysis was performed using edgeR (FDR < 0.05). **(C)** Bars represent the distribution of the 64 NaBut × Bel interaction-responsive genes across MapMan functional categories at 2 h. Functional categories represented by at least two genes are shown individually, whereas genes belonging to categories represented by a single gene or lacking a MapMan functional annotation were grouped as “Other / unassigned”. Red and blue bars indicate positive and negative interaction effects, respectively, reflecting shifts toward higher or lower expression in the presence of NaBut. Differential expression analysis was performed using edgeR (FDR < 0.05).

To determine whether this apparent modulation could be statistically supported at the gene level, a contrast-based analysis was used to identify genes exhibiting a significant NaBut × Belvine^®^ interaction (Fig. 6B, Fig. S4A). At 2 h, 1,085 DEGs were identified in response to Belvine^®^ alone, compared with 2,293 in response to Belvine^®^ in the presence of NaBut, showing that substantially more genes were differentially expressed when NaBut was present. Beyond this overall difference, the interaction contrast identified 64 genes for which the response to Belvine^®^ was significantly different depending on the presence of NaBut (Fig. 6B). Among the 64 genes, 21 showed a positive interaction effect, indicating that the response to Belvine^®^ shifted toward more positive values in the presence of NaBut (e.g., stronger induction or weaker repression), whereas 43 showed a negative interaction effect, indicating a shift toward more negative values (e.g., stronger repression or weaker induction). Of these 64 genes, 41 were assigned to MapMan functional categories (Fig. 6C, Table S6). Positive interaction genes were mainly associated with transport and metabolic processes, whereas negative interaction genes were predominantly associated with protein biosynthesis and RNA-related functions. Together, these results provide statistical support at the gene level for the NaBut-dependent modulation of the transcriptional response to Belvine^®^ suggested by the global analysis. The temporal persistence of this interaction was then examined at 4 h. The number of DEGs identified in response to Belvine^®^ increased in both the absence and presence of NaBut, reaching 2,950 and 5,495 genes, respectively, compared with 1,085 and 2,293 genes at 2 h. Despite this larger transcriptional response, only four genes showed a significant NaBut × Belvine^®^ interaction at 4 h (Fig. S4A), compared with 64 at 2 h, indicating that the effect of NaBut on the Belvine^®^ transcriptional response was predominantly detected at the earlier time point. Among the four genes, only one could be assigned to a MapMan functional category, precluding meaningful functional interpretation at this time point (Fig. S4B, Table S6).

### 3.8. Targeted transcriptomic analysis confirmed the synergy between NaBut and Belvine^®^ in the vineyard conditions

To assess whether the transcriptional modulation observed under controlled conditions was also detectable under vineyard conditions, the expression of a panel of defense-related genes was analyzed in leaves collected from an experimental vineyard following treatments with copper (Champ^®^ Flo Ampli, Nufarm) associated to Belvine^®^, with or without NaBut (Fig. S5A). These targeted expression data were then compared with the RNA-seq results obtained under controlled conditions. Despite differences in experimental conditions and sampling kinetics between both experiments, concordance in the direction of the NaBut effect was observed, with 671 of the 960 RNA-seq-vineyard comparisons (69.9%) showing the same direction of modulation (Fig. S5B; Table S3). Concordant responses were observed among genes associated with several defense-related processes, including hormone-related responses, PR proteins and secondary metabolism (Fig. S5C). Notably, NaBut shifted the Belvine^®^-associated expression response of several PAL and STS genes toward higher values in the RNA-seq dataset, and similar positive effects were repeatedly observed for the corresponding markers across vineyard sampling points (Table S3). Concordant positive effects were also observed for LOX9 and PR2/β-1,3-glucanase. Conversely, some genes showed concordant negative effects, indicating that NaBut-mediated modulation was not uniformly associated with increased gene expression. Overall, these results indicate that several transcriptional effects of NaBut on the response to Belvine^®^ identified under controlled conditions were similarly detected under vineyard conditions.

### 3.9. The disinhibition strategy enhances protection against *P. viticola* under vineyard conditions

Given that NaBut modulates the expression of Belvine^®^-induced defense genes under vineyard conditions, we sought to determine whether the NaBut-Belvine^®^ combination could enhance protection compared with Belvine^®^ alone, using two complementary experimental approaches. As NaBut alone did not confer direct protection under controlled conditions, these experiments specifically assessed the potential benefit of this strategy within a reduced-input program combining reduced copper application and Belvine^®^.

Because natural *P. viticola* infection is highly dependent on weather conditions and can vary markedly between growing seasons, a combined field-laboratory approach was first implemented in an experimental vineyard at Marsannay-la-Côte (Côte-d’Or, France). Treatments were applied to Chardonnay plants and leaves were subsequently collected 48 h later for controlled *P. viticola* inoculation in the laboratory. Bayesian model-adjusted estimates confirmed a lower leaf DM severity following reduced copper + Belvine^®^ + NaBut than following reduced copper + Belvine^®^ alone (Fig. 7A), with a probability of improved protection of 95.3%.

**Figure 7.**
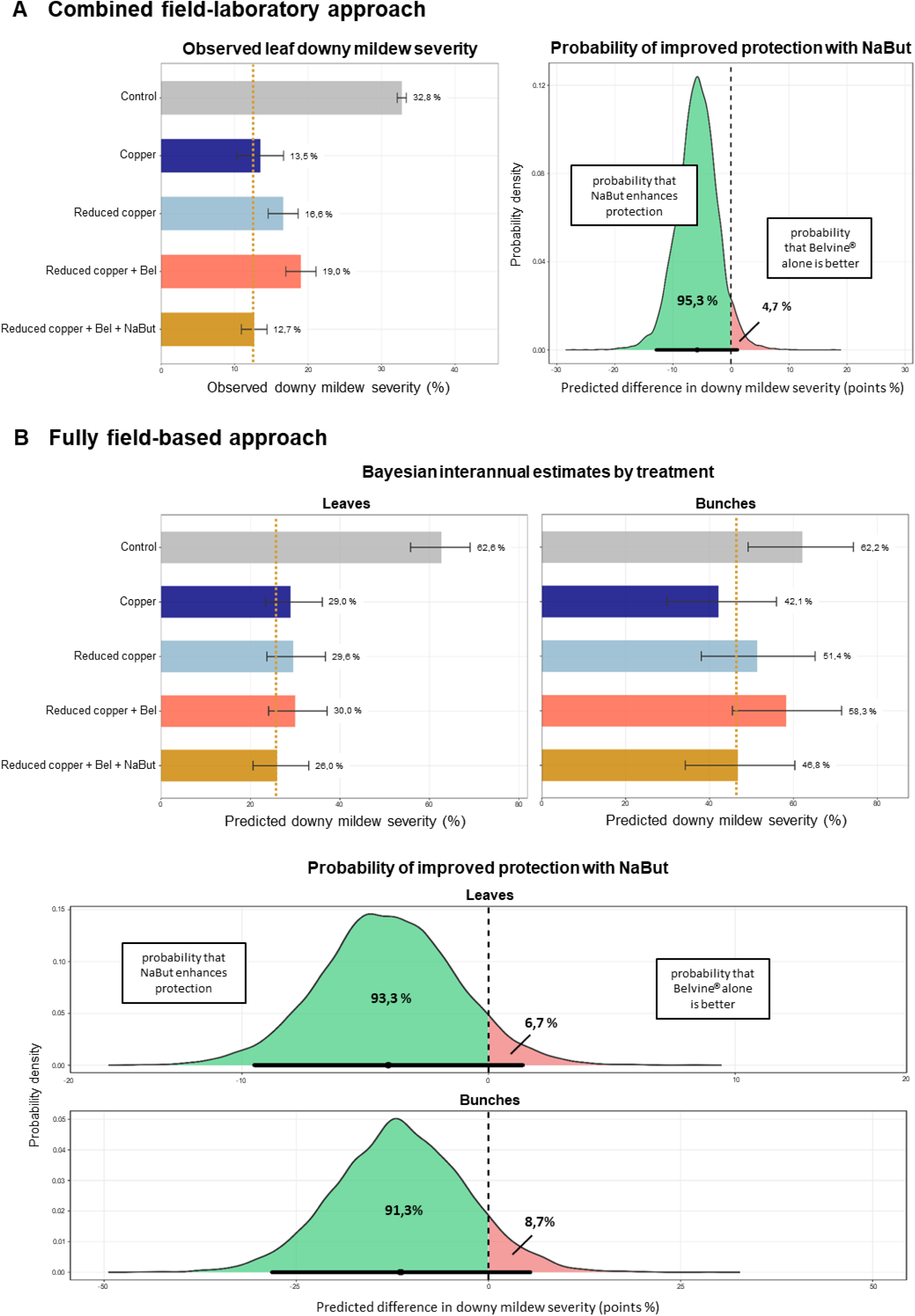
NaBut addition to Belvine^®^ is associated with improved grapevine protection against downy mildew under vineyard conditions. Experiments were conducted at two vineyard sites located in Marsannay-la-Côte (Côte-d’Or, France) and Azé (Saône-et-Loire, France), using two complementary experimental approaches. At Marsannay-la-Côte, treated leaves were collected 48 h after treatment from the vineyard and subsequently inoculated with *P. viticola* under laboratory conditions. At Azé, treatments were evaluated directly in the vineyard over two growing seasons, following artificial *P. viticola* inoculation and misting to favour disease development. Treatments included a water-treated control, copper applied at the full (Copper) or half (Reduced copper) annual field rate, reduced copper combined with Belvine**^®^** at half rate (Reduced copper + Bel), and the same treatment supplemented with NaBut (Reduced copper + Bel + NaBut). The full rate corresponded to 3.9 kg/ha for copper and 10 mL.L^-1^ for Belvine**^®^**. NaBut was applied at 220 mg.L^-1^ (2 mM) **(A)** Combined field-laboratory approach (Marsannay-La-Côte). Observed leaf downy mildew severity for each treatment is shown as the mean ± SEM of three independent experiments (left). Posterior distribution of the difference in predicted severity between Reduced copper + Bel + NaBut and Reduced copper + Bel (right). **(B)** Fully field-based approach (Azé). Bayesian model-adjusted estimates of downy mildew severity for leaves and bunches across the two consecutive growing seasons (top), and corresponding posterior distributions of the difference in predicted severity between Reduced copper + Bel + NaBut and Reduced copper + Bel for leaves and bunches (bottom). Treatment and year were included as additive effects to account for interannual variation, and posterior predictions were averaged with equal weight across the two growing seasons. Bars represent posterior means and error bars indicate 95% credible intervals. In the posterior distribution plots, the x-axis represents the predicted difference in disease severity in percentage points, with negative values indicating improved protection following NaBut addition. Green and red areas represent the posterior probabilities of improved protection (difference < 0) and no improvement (difference > 0), respectively. Points and horizontal lines indicate posterior mean differences and their 95% credible intervals.

Following these results, the strategy was further evaluated under fully field-based conditions in a commercial Chardonnay vineyard at Azé (Saône-et-Loire, France). The trial was conducted over two consecutive growing seasons, and DM severity was assessed on both leaves and bunches. Bayesian interannual estimates showed lower DM severity following reduced copper + Belvine^®^ + NaBut than following reduced copper + Belvine^®^ alone on both leaves and bunches (Fig. 7B), with probabilities of improved protection of 93.3% and 91.3%, respectively. Across both experimental approaches and plant organs, probabilities consistently supported a beneficial effect of NaBut supplementation within the reduced copper + Belvine^®^ program, supporting the relevance of the disinhibition strategy under vineyard conditions.

Moreover, in the context of the French Ecophyto 2030 strategy, which aims to reduce the use and risk of plant protection products by 2030, model-adjusted severity under reduced copper + Belvine^®^ + NaBut was also compared with that under the full copper application rate at both sites. At Marsannay-la-Côte, DM severity under reduced copper rate + Belvine^®^ + NaBut (12.7%) was numerically similar to that under the full copper application rate (13.5%) and lower than that under reduced copper alone (16.6%; Fig. 7A). At Azé, model-adjusted severity estimates showed a similar pattern on leaves, with lower severity under reduced copper + Belvine^®^ + NaBut than under both full-rate copper (26.0% vs 29.0%) and reduced copper alone (26.0% vs 29.6%; Fig. 7B). On bunches, severity under reduced copper + Belvine^®^ + NaBut (46.8%) was intermediate between that under the full-rate copper (42.1%) and reduced copper alone (51.4%). Overall, the reduced-copper program supplemented with Belvine^®^ and NaBut provided disease severity estimates that were similar to or lower than those obtained with the full-rate copper in several comparisons, while showing lower severity than reduced copper alone. Altogether, these findings support the field applicability of the disinhibition strategy and its potential to maintain effective disease protection while reducing conventional crop protection inputs.

## Discussion

The use of inhibitors targeting negative regulators of grapevine immunity remains poorly studied. We therefore focused on the HD2 HDAC-subfamily, whose members have been identified as negative regulators of immunity in rice and tobacco (Bourque et al., 2011; Ding et al., 2012). Two HD2 proteins, VvHD2a and VvHD2b, have been identified in grapevine and share conserved domains, including a nuclear localization signal, with the rice protein OsHDT701 (Ding et al., 2012) and tobacco NtHD2a / NtHD2b (Bourque et al., 2016).

Using *A. thaliana* as a heterologous model, we showed that constitutive expression of VvHD2a or VvHD2b increased susceptibility to the necrotrophic fungus *B. cinerea*. This result is consistent with previous observations in rice, where overexpression of the ortholog OsHDT701 increased susceptibility to fungal infection (Ding et al., 2012). Similarly, silencing of NtHD2a and NtHD2b in tobacco enhanced defense responses following treatment with a PDI, namely cryptogein of the oomycete *Phytophtora cryptogea* (Bourque et al., 2011). Together, these observations support a conserved negative role for HD2 proteins across plant species, at least in the context of fungal or oomycete-associated defense responses. This is reinforced by the fact that HD2s are highly conserved across plant species (Bourque et al., 2016).

To investigate the mechanisms underlying the increased susceptibility of *A. thaliana VvHD2a/b* overexpressing lines to *B. cinerea*, we examined PTI-associated responses. Overexpression of VvHD2a or VvHD2b reduced flg22-induced ROS production and repressed the expression of defense-related genes involved in JA and indole glucosinolate (IGS) pathways. The repression of JA-associated genes is particularly relevant because JA signaling plays an important role in resistance to *B. cinerea* (Smirnova et al., 2017; Yang et al., 2007). Consistent with these observations, *AtLOX4* expression induced with flg22 treatment was reduced in VvHD2a- and VvHD2b-overexpressing plants, which were also more susceptible to *B. cinerea*. Although our results do not demonstrate direct regulation of *AtLOX4* by VvHD2 proteins, they support a potential role for VvHD2a and VvHD2b in restricting JA-associated defense responses. We also observed reduced expression of two genes involved in IGS precursor biosynthesis in VvHD2a- and VvHD2b-overexpressing plants. IGS-derived metabolites contribute to plant defense against several pathogens, including *B. cinerea* (Bednarek et al., 2009; Buxdorf et al., 2013; Clay et al., 2009; Kliebenstein et al., 2005). Recent studies further showed that histone modifications can regulate glucosinolate biosynthesis during stress responses in *Arabidopsis* (Choi et al., 2026), while HDA9 has been reported as being implicated in glucosinolate regulation in other contexts (Choi et al., 2024). Therefore, our results raise the possibility that VvHD2 proteins contribute to the repression of the IGS pathway during plant responses to pathogen. Overall, our results support a model in which VvHD2a and VvHD2b negatively regulate specific immune responses, including flg22-induced ROS production and the activation of JA- and IGS-associated defense pathways in *A. thaliana*. Their overexpression is associated with increased susceptibility to *B. cinerea*. Conversely, it is associated with resistance to *P. syringae* as previously demonstrated in *A. thaliana,* thus highlighting a pathogen-dependent function of HD2 proteins that would merit to be further studied (Latrasse et al., 2017).

We next investigated whether inhibition of HDAC could alleviate this negative regulation. Treatment with NaBut, a well-known HDAC inhibitor (Steliou et al., 2012; Xiong et al., 2016), significantly reduced the enhanced susceptibility of *VvHD2a*- and *VvHD2b*-overexpressing plants to *B. cinerea*, whereas no comparable effect was observed in WT plants under our experimental conditions. These results suggest that HDAC inhibition could partially counteract the increased defense repression associated with *VvHD2* overexpression. Although NaBut is not specific to HD2 proteins and can affect multiple HDACs (Deng et al., 2025), these findings provide proof of concept that targeting HDAC may help enhance defense responses in crops. Furthermore, combining HDAC inhibition with a PDI, as demonstrated in tobacco by Bourque et al. (2011), could represent a complementary strategy to enhance plant immunity and potentially improve the effectiveness of PDI.

This strategy was tested using grapevine. We combined NaBut with Belvine^®^ (a PDI patented for DM and PM in grapevine) and evaluated their effects against *P. viticola*. Our results showed that NaBut alone did not confer protection, whereas its combination with a reduced dose of Belvine^®^ significantly increased protection, revealing a synergistic interaction between the two treatments. This suggests that HDAC inhibition is not sufficient on its own to establish effective resistance against *P. viticola*, but can potentiate defense responses once they have been induced. A similar link between HDAC inhibition and enhanced plant immunity has previously been reported in rice. Xu et al. (2022) showed that NaBut treatment enhanced resistance to *Magnaporthe oryzae*, together with increased H3K9 acetylation and expression of several defense-related genes. However, the authors also found that other HDAC inhibitors increased global histone acetylation without improving resistance, indicating that increased histone acetylation alone is not necessarily sufficient to confer protection. Our results extend this concept by suggesting that HDAC inhibition can potentiate the response triggered by a PDI. This concept is consistent with previous studies showing that other compounds such as acibenzolar-S-methyl and sulforaphane, can promote chromatin modifications associated with enhanced transcriptional activation of defense-related genes upon subsequent stimulation (Jaskiewicz et al., 2011; Schillheim et al., 2018). In our study, the transcriptomic data provide further support for this model. Together, our observations suggest that Belvine^®^ may shape the direction of the transcriptional response, while NaBut appears to modulate its magnitude. This modulation notably affects defense-associated events, including ROS production, hormone signaling and lignin-related responses, indicating that the effect of NaBut extends to pathways relevant to plant immunity. More importantly, the interaction analysis demonstrates that the transcriptional response to Belvine^®^ itself is modified by the presence of NaBut. After 2 h of treatment, 64 genes showed a significant NaBut × Belvine^®^ interaction, indicating that their response to Belvine^®^ differed depending on whether NaBut was used. Both positive and negative interactions were detected, showing that NaBut does not simply enhance Belvine^®^-induced transcription, but rather modulates the way in which specific genes respond to the PDI. The marked decrease from 64 significant interactions after 2 h of treatment to only four after 4 h further indicates that this effect is predominantly associated with the early establishment of the Belvine^®^ response. This rapid and transient effect is particularly interesting in light of previous work on NaBut-mediated HDAC inhibition. In mammalian cells, Wang et al. (2018) reported detectable changes in the histone acetylome as early as 10 min after NaBut treatment, demonstrating that NaBut-induced changes in histone acetylation can occur rapidly. In plants, Xu et al. (2022) showed that NaBut-induced changes in the expression of several defense-related genes were transient in rice, whereas H3K9 acetylation at these loci remained in a sustained state. The authors suggested that additional regulatory mechanisms may subsequently constrain gene expression despite the maintenance of an acetylated chromatin state. Together, these findings suggest that chromatin modifications may contribute to a primed state that facilitates subsequent defense gene regulation, without necessarily sustaining constitutive gene expression. Although our study does not directly assess histone acetylation at the genes affected by the NaBut × Belvine^®^ interaction, the predominantly early transcriptional effect observed here supports the idea that HDAC inhibition may influence a regulatory window during the establishment of the defense response. We therefore propose that Belvine^®^ provides the defense-inducing signal, while NaBut modifies the early transcriptional response to this signal. In this model of disinhibition and activation of plant immune response, HDAC inhibition would not generate an independent defense program, but would relieve regulatory constraints that shape the response triggered by Belvine^®^, thereby contributing to its potentiation. An important next step was to determine whether the benefit of this strategy could be maintained under vineyard conditions. Indeed, although PDIs can provide effective protection under controlled conditions, their efficacy in vineyards remains variable and can be influenced by multiple biotic and abiotic factors (Delaunois et al., 2014; Héloir et al., 2019). Based on the synergistic effect observed between NaBut and Belvine^®^ under controlled conditions, we therefore incorporated NaBut into reduced-input protection programs combining Belvine^®^ with reduced copper rates. The benefit of NaBut addition was first supported in an experimental vineyard using a combined field-laboratory approach, with a 95.3% probability of improved protection. This benefit was further supported under fully field-based conditions in a vineyard under real-world production conditions, with probabilities above 90% of improved protection on both leaves and bunches across two consecutive growing seasons. The integration of PDI into reduced-input vineyard protection programs has already shown promising results. Taibi et al. (2023), for example, demonstrated over three growing seasons that resistance inducers could contribute to the control of grapevine DM and PM when combined with low doses of copper and sulfur. Similarly, Piancatelli et al. (2024) showed that resistance-inducing treatments combined with reduced copper inputs could, under some conditions, provide disease control comparable to reference copper-based programs. In addition to this variability in field efficacy, the economic cost of plant defense inducers may represent another constraint, particularly when repeated applications or substantial product doses are required. The high production cost of some grapevine PDIs has indeed been identified as a potential barrier to their transfer to vineyard applications (Jeandet et al., 2023). Improving PDI efficacy while reducing the amount of product required therefore represents an important objective for their practical implementation. Our strategy addresses this issue from a different perspective by combining molecules with complementary modes of action: defense induction and defense disinhibition. Rather than increasing the amount of PDI applied, NaBut was used to enhance the response induced by a reduced dose of Belvine^®^. Under our vineyard programs, Belvine^®^ was applied at half or one-quarter of the recommended field rate, depending on the experimental site, together with a 50% reduction in copper input. Despite these substantial reductions, disease levels obtained with the combined NaBut × Belvine^®^ strategy were close to those observed with the full-rate copper reference and were numerically lower in some comparisons. Thus, the benefit of NaBut was not limited to improving protection within a reduced-input program: defense disinhibition allowed a reduced dose of PDI, combined with reduced copper, to approach the level of disease control achieved with full-rate copper. The convergence of results obtained under controlled conditions, transcriptomic analyses and vineyard experiments supports defense disinhibition as a promising strategy to potentiate PDI-based crop protection, at least in grapevine, while reducing both PDI and fungicide inputs.

## Conclusion

To conclude, our findings show that effective plant protection depends not only on activating defense responses but also on relieving the mechanisms that restrain them. Combining a PDI with a HDAC inhibitor potentiated an otherwise limited PDI-induced defense response, with the PDI providing the defense signal and HDAC inhibition enhancing its early transcriptional effects. This approach could mitigate the limitations associated with the variable efficacy and cost of defense inducers by allowing their application at lower doses in combination with low-cost NaBut, thereby providing a complementary route toward more sustainable crop protection. Further work on the underlying epigenetic mechanisms and on robustness across environments and cultivars will be needed.

## APPENDIX SUPPLEMENTARY DATA

Figure S1. Alignement and structural features of type-2 HDACs from Vitis vinifera

Figure S2. Bacterial quantification after inoculation (Day 0) of A. thaliana with Pseudomonas syringae pv. tomato DC3000

Figure S3. Effects of NaBut and Belvine® on B. cinerea mycelial growth and P. viticola zoospore motility at concentrations used for plant treatments

Figure S4. NaBut × Belvine® transcriptional interaction at 4 h

Figure S5. Gene expression responses to Belvine® and NaBut under vineyard conditions and comparison with transcriptional responses observed under controlled conditions

Table S1. Culture media for in vitro grapevine plantlets

Table S2. Primer list

Table S3. Concordance responses

Table S4. HDAC sequences

Table S5. RNAses Mapman functional analysis

Table S6. RNAseq gene interactions

Video S1. Control treatment

Video S2. NaBut 0.1 mM treatment

Video S3. NaBut 1 mM treatment

Video S4. NaBut 5 mM treatment

Video S5. NaBut 10 mM treatment

Video S6. NaBut 1 mM – Belvine 0.75 g.L^-1^ treatment

Video S7. Belvine 0.75 g.L^-1^ treatment

Video S8. Belvine 3.25 g.L^-1^ treatment

## ACKNOWLEDGEMENTS

The authors acknowledge the support of the French National Research Agency (ANR) under the grants 20-PCPA-0010 (PPR Vitae, Cultivating the grapevine without pesticides: towards agroecological wine-producing socio-ecosystems) and ANR-23-ECOM-0008 (Ecophyto Maturation, Derepression of plant imnune responses: a new strategy to improve biocontol efficiency). We also thank Chrystel Deulvot for confocal microscopy support from the regional Centre of Microscopy/ DImaCell platform (Dijon, France).

## AUTHOR CONTRIBUTIONS

Jérémy Villette: Conceptualization, Formal analysis, Investigation, Writing – original draft Chloé Ilbert: Conceptualization, Formal analysis, Investigation, Software, Writing – original draft

Sébastien Aimé: Investigation

Manon Chargy: Investigation

Marie-Cécile Dufour: Investigation

Cécile Blanchard: Investigation

Caroline Bonneau: Methodology, Investigation

Karine Palavioux: Investigation

Hoai-Nam Truong-Cellier: Investigation

Nathalie Harzic: Writing – review and editing

Marielle Adrian: Writing – review and editing

Stephane Bourque: Conceptualization, Funding acquisition, Investigation, Supervision, Writing – review and editing

## DATA AVAILABILITY

All data supporting the findings of this study are available within the paper and its supporting information are published online, and the materials of this study are available from the corresponding author upon reasonable request.

