## Supplementary material for "Disinhibition of Plant Defense Responses as an Innovative Biocontrol Strategy in Grapevine": Figure S1

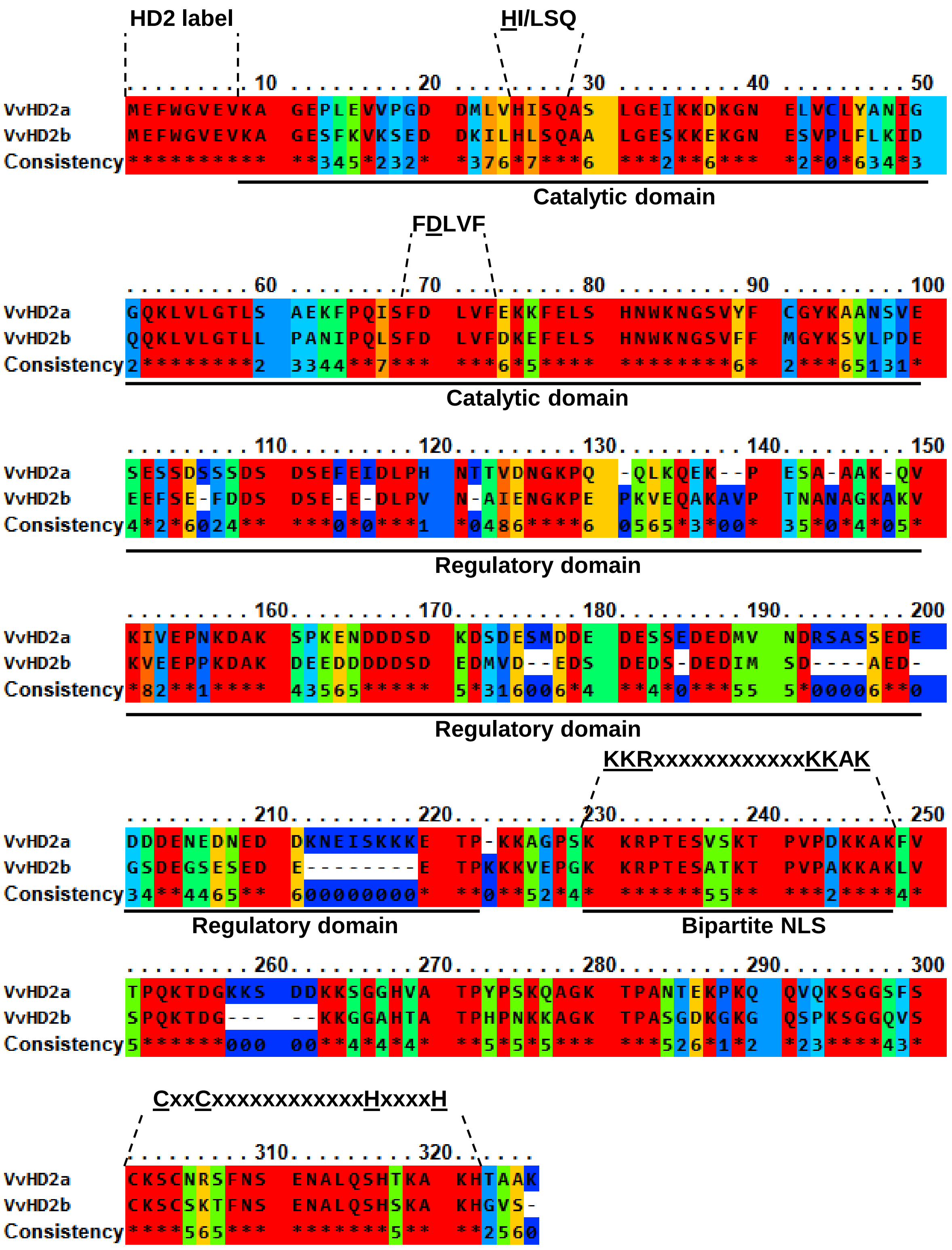


**Figure S1. Alignement and structural features of type-2 HDACs from *Vitis vinifera*.** Alignment of VvHD2a (320 aa) and VvHD2b (300 aa) proteins reveals 69.6% sequence similarity, with conservation of well-characterized motifs and domains. Both VvHD2a and VvHD2b contain the structural feature specific to the HD2 family (HD2 label). They also possess the conserved residues H25 and D70, which are important for catalytic activity and are located within the predicted catalytic domain spanning residues 1-99. Both proteins further contain a divergent regulatory domain and a highly conserved bipartite nuclear localization signal (NLS), with the corresponding motif underlined. Finally, both grapevine HD2 proteins contain a C-terminal zinc-finger domain characterized by conserved motifs, which are underlined. Domain and motif predictions were performed using InterProScan and based on the features described by Bourque et al. (2016).
