## Supplementary material for "Disinhibition of Plant Defense Responses as an Innovative Biocontrol Strategy in Grapevine": Figure S2

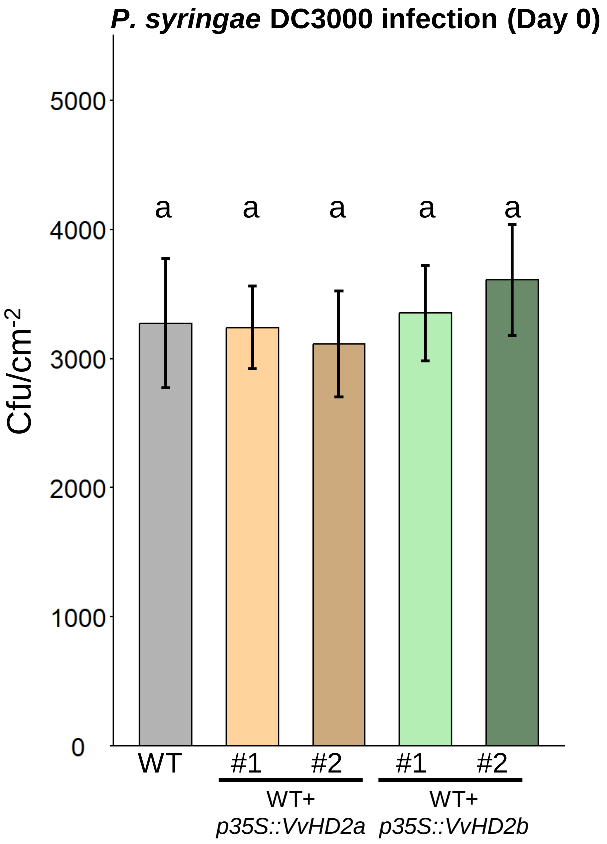


**Figure S2. Bacterial quantification after inoculation (Day 0) of *A. thaliana* with *Pseudomonas syringae* pv. *tomato* DC3000**. Bars represent the mean (± SE) colony-forming units (CFU) per cm² of leaf area in WT plants (grey), WT+*p35S::VvHD2a#1,#2 (*light and dark orange) and WT+*p35S::VvHD2b#1,#2* (light and dark green), measured from three independent biological experiments. Letters indicate no statistically significant differences among genotypes (Kruskal–Wallis test followed by Dunn's multiple comparison test with Bonferroni correction, $P < 0.05$).
