## Supplementary material for "Disinhibition of Plant Defense Responses as an Innovative Biocontrol Strategy in Grapevine": Figure S3

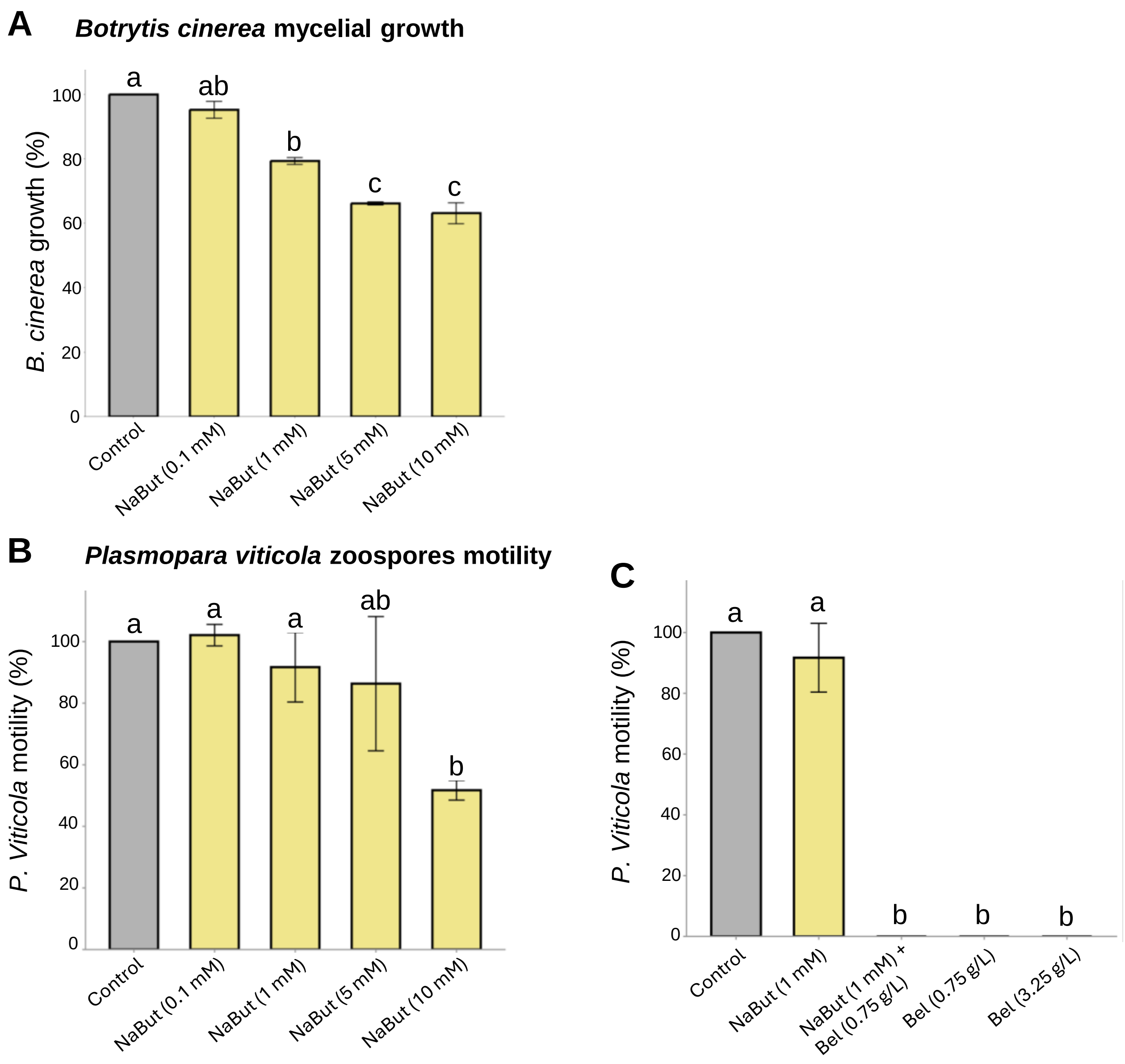


**Figure S3. Effects of NaBut and Belvine® on *B. cinerea* mycelial growth and *P. viticola* zoospore motility at concentrations used for plant treatments.** (A) Direct effect of NaBut at increasing concentrations (0.1, 1, 5, and 10 mM) on *B. cinerea* mycelial growth. Conidia were directly exposed to the indicated concentrations, and fungal growth was quantified after 60 h and expressed relative to the $H_2O$-treated control. Bars represent means ± SD of three independent biological experiments, each comprising three technical replicates. Different letters indicate significant differences among treatments (Kruskal-Wallis test followed by Dunn's multiple-comparison test with Holm's correction, $P < 0.05$). (B) Dose-dependent effect of NaBut on *P. viticola* zoospore motility at increasing concentrations (0.1, 1, 5, and 10 mM). (C) Direct effects of NaBut and Belvine®, applied alone or in combination, on *P. viticola* zoospore motility. For panels B and C, zoospores were directly exposed to the indicated treatments, and motility was expressed relative to the $H_2O$-treated control. Bars represent means ± SD of three independent biological experiments, each comprising six technical replicates. Different letters indicate significant differences among treatments (one-way ANOVA followed by Tukey's HSD test, $P < 0.05$).
