## Supplementary material for "Disinhibition of Plant Defense Responses as an Innovative Biocontrol Strategy in Grapevine": Figure S4

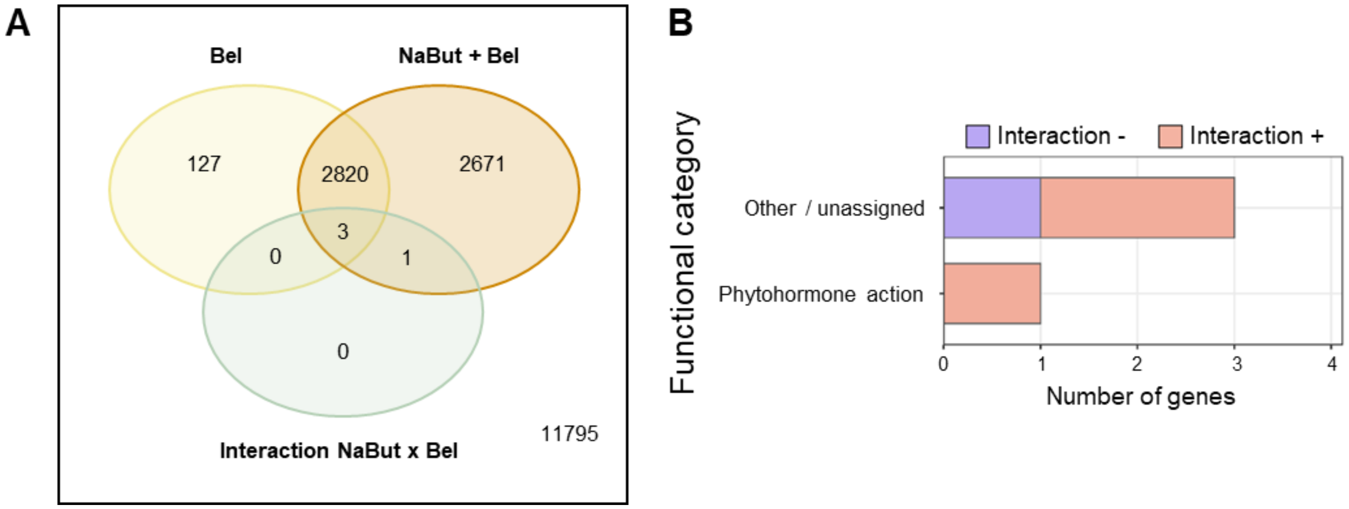


**Figure S4. NaBut × Belvine® transcriptional interaction at 4 h. (A)** Venn diagram showing the number of DEGs identified at 4 h in response to Belvine® (Bel) alone, the Belvine® effect in the presence of NaBut (NaBut + Bel), and genes displaying a significant NaBut × Bel interaction effect. Differential expression analysis was performed using edgeR (FDR < 0.05). Numbers indicate the count of significant genes in each contrast and their overlaps. **(B)** Distribution of the four genes displaying a significant NaBut × Bel interaction effect at 4 h according to MapMan functional categories. Genes lacking a MapMan functional annotation were grouped as "Other / unassigned". Red and blue bars indicate positive and negative interaction effects, respectively, corresponding to shifts toward higher or lower expression in the presence of NaBut. Differential expression analysis was performed using edgeR (FDR < 0.05).
