## Supplementary material for "Disinhibition of Plant Defense Responses as an Innovative Biocontrol Strategy in Grapevine": Figure S5

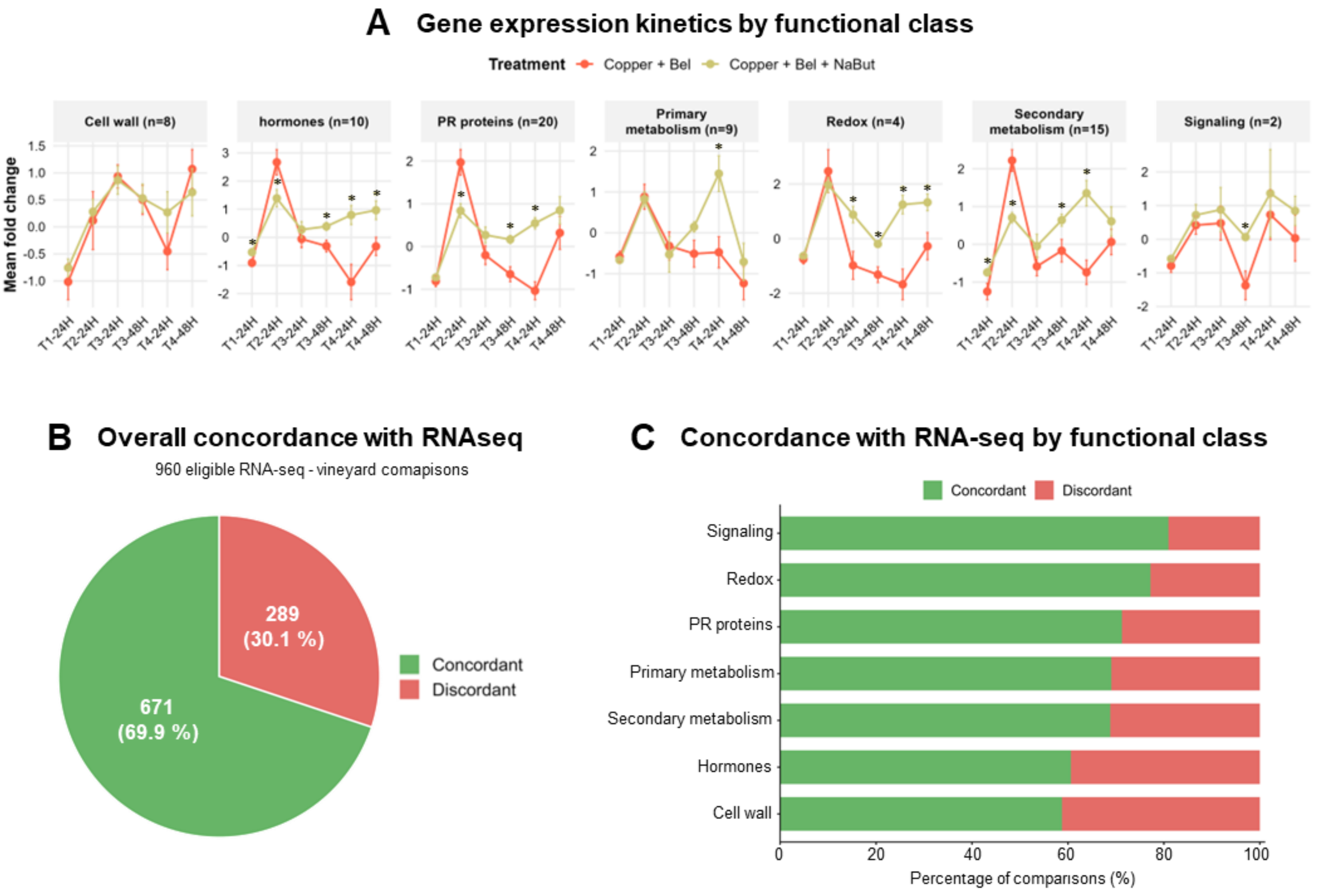


**Figure S5. Gene expression responses to Belvine® and NaBut under vineyard conditions and comparison with transcriptional responses observed under controlled conditions. (A)** Gene expression kinetics in *Vitis vinifera* cv. Cabernet Sauvignon leaves collected from an experimental vineyard in Bordeaux (Gironde, France). Leaves were sampled 24 or 48 h after successive treatments (T1 to T4, corresponding to treatments 1 to 4). Gene expression was analyzed using a high-throughput microfluidic qPCR approach based on the NeoViGen96/BioStim96 gene panels described by Dufour et al. (2016) and Bodin et al. (2020). Genes were grouped into functional classes, and expression is presented as the mean fold change ± SEM for each class. The genes included in each functional class are reported in Table S5. Copper + Bel and Copper + Bel + NaBut treatments were compared at each sampling time. Asterisks indicate significant differences between the two treatments (t-test followed by Benjamini-Hochberg correction, adjusted $p < 0.1$). **(B)** Overall concordance in the direction of the NaBut effect between the RNA-seq dataset obtained under controlled conditions and the vineyard dataset. After matching genes from the vineyard panel with corresponding Vitvi genes and applying the RNA-seq significance criteria independently at 2 h and 4 h, 160 eligible gene associations were retained across the two RNA-seq time points. Each was compared with the six vineyard sampling times, resulting in 960 comparisons. **(C)** Distribution of concordant and discordant RNA-seq–vineyard comparisons within each functional class. Concordance was defined as the same direction of the NaBut effect in both datasets, based on the sign of Delta RNA-seq and Delta Vineyard, whereas comparisons with opposite signs were classified as discordant. Complete comparison data and the calculation of Delta RNA-seq and Delta Vineyard are provided in Table S2.
